# A Microstructurally-Motivated Framework to Study Autoregulation in the Coronary Circulation

**DOI:** 10.64898/2025.12.02.691683

**Authors:** Matthew J. Eden, Hamidreza Gharahi, Victoria E. Sturgess, Domingo E. Uceda, Seungik Baek, Daniel A. Beard, Johnathan D. Tune, C. Alberto Figueroa

## Abstract

Coronary autoregulation maintains relatively constant myocardial flow over a wide range of perfusion pressures through myogenic, shear-dependent, and metabolic control mechanisms. Understanding this phenomenon is challenging due to the coupled nature of these mechanisms and their heterogeneous effects throughout the coronary tree. In this study we developed a novel microstructurally-motivated model of coronary autoregulation based on constrained mixture theory, with anatomical and structural parameters calibrated through a homeostatic optimization framework. Autoregulation was simulated at three myocardial depths (subepicardium, midwall, and subendocardium), with the calibrated model accurately reproducing baseline hemodynamics and autoregulatory responses. For changes in epicardial pressure, our model reproduced experimentally measured subendocardium-to-subepicardium flow ratios (ENDO/EPI) and changes in vessel diameter, demonstrating its predictive capability. Furthermore, we extended Womersley’s theory to simulate phasic coronary hemodynamics with a time-varying intramyocardial pressure. This microstructurally-motivated framework provides a mechanistic foundation for investigating coronary autoregulation and long-term vascular growth and remodeling in pathphysiological conditions.

**Summary:**

- Coronary autoregulation is defined as the capability of the coronary circulation to maintain the blood supply to the heart over a range of perfusion pressures. This phenomenon is facilitated through intrinsic mechanisms that control the vascular resistance by regulating the mechanical function of smooth muscle cells. Understanding the mechanisms involved in coronary autoregulation is one of the most fundamental questions in coronary physiology.
- This paper presents a structurally-motivated coronary autoregulation model that uses a nonlinear continuum mechanics approach to account for the morphometry and vessel wall composition in two coronary trees in the subepicardial and subendocardial layers.
- The model is calibrated against diverse experimental data from literature and is used to study heterogeneous autoregulatory response in the coronary trees. This model drastically differs from previous models, which relied on lumped parameter model formulations, and is suited to the study of long-term pathophysiological growth and remodeling phenomena in coronary vessels.

## 1. INTRODUCTION

The myocardium extracts approximately 70-80% of oxygen from coronary arterial blood under resting conditions. Therefore, shifts in metabolic demand (myocardial oxygen consumption, MVO_2_) are largely met by changes in coronary blood flow [1]. Coronary flow regulation is the result of vessel diameter modulation via local control of the active stress in the vascular smooth muscle cells (SMCs) [2]. One aspect of coronary flow regulation is referred to as pressure-flow autoregulation, in which changes in coronary perfusion pressure are met with adjustments in microvascular resistance to maintain a relatively constant coronary blood flow at a given MVO_2_.

Coronary autoregulation relies on SMCs reactivity, regulated through myogenic, metabolic, and shear-dependent control mechanisms. Myogenic control is the intrinsic response of SMCs to changes in local wall stress [3]. Metabolic control is a local feedback mechanism where either an increase in local oxygen consumption or oxygen extraction from the myocardial blood supply triggers a vasodilatory signal [1]. Shear-dependent control is a dilatory mechanism mediated via shear-induced production of nitric oxide (NO) by the endothelial cells. Studies have shown that the relative strengths of these control mechanisms vary throughout the coronary circulation [4]. Myogenic control has been observed to be small in precapillary arterioles and large in 50-150 μm diameter arteries [5]. Shear-dependent control is most active in the arteries and large arterioles [6]. Lastly, the signal for the metabolic feedback mechanism is initiated in the capillaries, conducted upstream, and gradually decays for vessels >150 μm diameter [7].

In additional to the variable strengths of the autoregulatory mechanisms, the coronary circulation contains several other important heterogeneities. Non-uniform myocardial-vessel interactions result in subendocardial vessels experiencing a greater intramyocardial pressure than those in the subepicardium [8]. Furthermore, vessel morphometry varies transmurally across the myocardium, with vessels of the same diameter being thicker in the subepicardium versus the subendocardium [9]. Collectively, anatomical and functional differences across the myocardium result in a transmural variation in coronary autoregulation [10]. The inherent heterogeneous nature of the coronary circulation, combined with the challenges associated with obtaining *in-vivo* measurements, have made our understanding of coronary autoregulation to be limited and mostly phenomenological. Developing computational models that integrate available experimental data on morphometry, biomechanical, and physiological responses can enhance our understanding of coronary autoregulation in health, disease, and treatment planning conditions.

Coronary autoregulation has been the subject of numerous modeling studies over the last 50-60 years. Virtually all such studies have relied on lumped-parameter (0D) approaches. Liao and Kuo [5] and Cornelissen et al. [11] investigated the interaction and balance between autoregulatory mechanisms in the coronary circulation. While these studies incorporated the heterogeneity of the microvascular response, they partitioned the coronary tree into 4 compartments and ignored the interactions between vessels and myocardium. Pradhan et al. [12] develop a data-driven closed loop model of regulation using *in-vivo* data on coronary flow and oxygen extraction in response to exercise-induced changes in demand and perfusion pressure. The Pradhan model, which represents parallel control pathways using a block-diagram approach, does not represent explicit spatial features. Namani and colleagues [13] developed a coronary microcirculation model that integrated dynamic effects of flow with myogenic, shear dependent, and metabolic feedback control mechanisms in subendocardial and subepicardial vessels. Their model considered simple pressure-diameter rules to define the vessel behavior. Recently, Gharahi et al. [14] and Sturgess et al. [15] utilized a nonlinear three-layer lumped parameter network to investigate autoregulation across the myocardium under varying metabolic demands. While all these studies provide reasonable descriptions of autoregulatory responses, their inherent 0D nature complicates their interpretation. 0D models make it difficult to incorporate microstructurally-derived information on structure and function on image-based networks of vessels. Lastly, given that the short-term regulation of SMC tone and the long-term vascular growth and remodeling processes, including pathophysiologic responses, are highly intertwined [16], we submit that there is a pressing need to develop microstructurally motivated models of coronary autoregulation.

Constrained mixture models (CMMs) have been widely applied to describe the nonlinear mechanical behavior of arterial tissue, accounting for the contributions of the main load-bearing constituents (e.g., collagen, elastin and SMCs) [17], [18], [19]. This theory provides a formal means to represent mechanical function of a vessel based on the properties and relative mass fractions of its constituents. Using this theory, long-term growth and remodeling is represented by changes in the composition of the vessel wall. Previous applications of CMMs in vascular mechanics have focused on large arteries where homeostatic stress has been assumed constant. However, wall stress has been found to be size-dependent through the vasculature [20], [21]. More recently, the application of CMMs have been applied to study flow-mediated growth and remodeling (Fluid-solid growth) in both large vessels [22], [23] as well as network of vessels [24], [25]. However, CMMs have yet to be used to study autoregulation in vascular networks.

The goal of this study is to develop a microstructurally-motivated computational framework of coronary autoregulation, leveraging a recent homeostatic optimization model based on a CMM developed by Gharahi et al. [24]. The framework provides a means to estimate baseline activation of SMCs and homeostatic stresses, while incorporating experimental data on pressure-diameter, tree morphometry, and pulsatile hemodynamics. The utility of the proposed microstructurally-motivated framework is demonstrated through different examples of short-term adaptations to changes in perfusion pressure. A key benefit of this framework is that it provides a method to probe relative contributions of the different autoregulatory mechanisms across different vessel sizes and myocardial depths.

## 2. METHODS

### 2.1. Framework overview and calibration strategy

**Figure 1A** shows an overview of the framework. Anatomically, a symmetrically bifurcating coronary tree model is defined at three different myocardial depths: subepicardium (subepi), midwall, and subendocardium (subendo), which are subjected to an intramyocardial pressure. Vessel wall mechanics are governed by a microstructurally-motivated model based on a CMM formulation, treating collagen, SMCs, and elastin as load-bearing constituents. SMC activation modulates vessel diameter, and is controlled by myogenic, metabolic, and shear-dependent autoregulatory stimuli and determined through hemodynamic modeling. Our framework is calibrated using extensive literature data and considers the relative contributions of the three autoregulatory stimuli.

**Figure 1:**
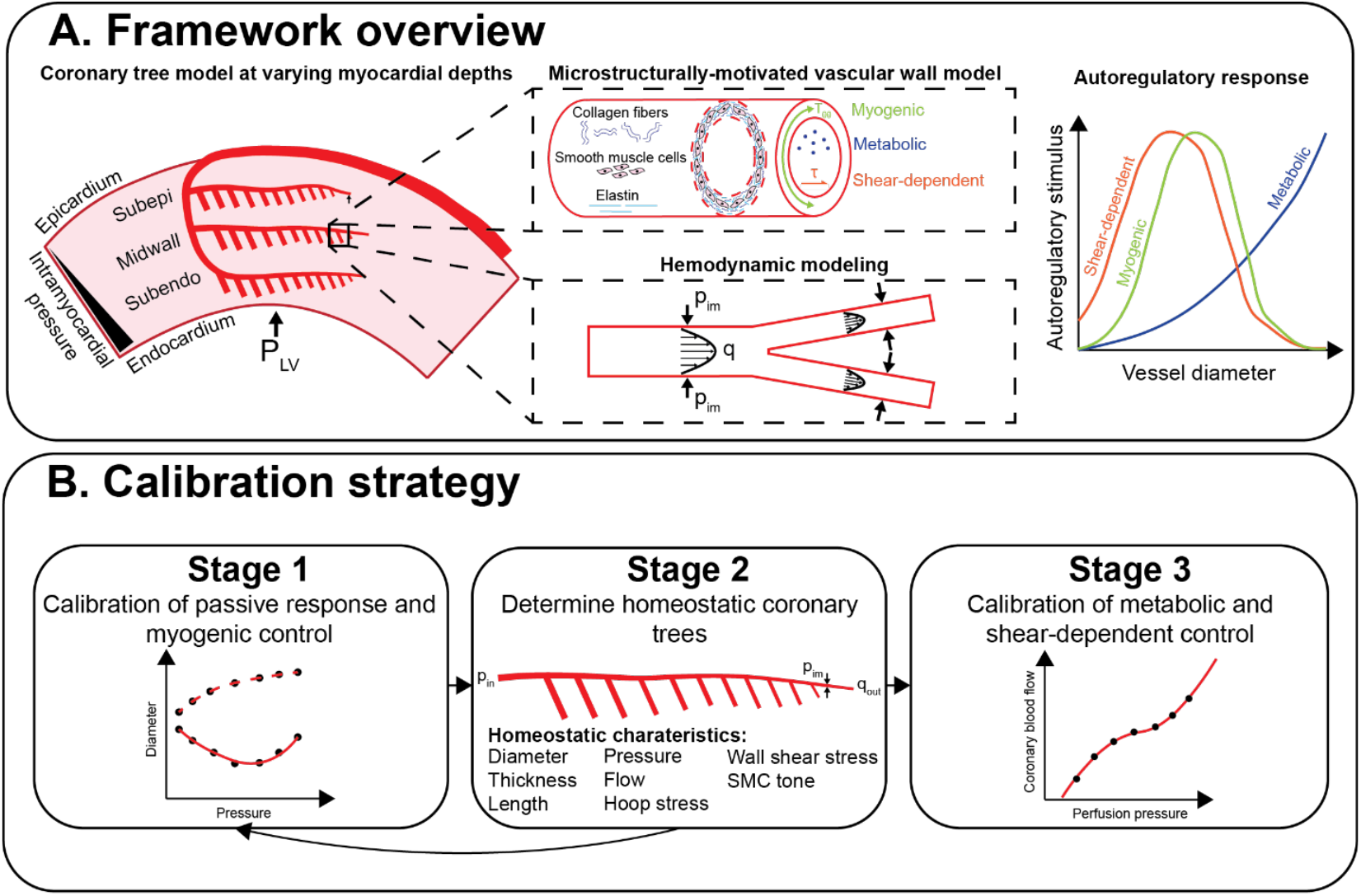
Overview of the coronary tree autoregulation framework (**A**), consisting of subtrees at three myocardial depths: subepi, midwall, and subendo subjected to spatially-varying intramyocardial pressure (*p*^*im*^). Vessel mechanics are governed by a microstructure-motivated wall model incorporating collagen, SMCs, and elastin. SMC activation is controlled by myogenic, metabolic, and shear-dependent autoregulatory stimuli, which are calculated through hemodynamic modeling. The model undergoes a three-stage calibration procedure (**B**) to establish passive and active properties that reproduce coronary pressure-flow autoregulation.

Framework calibration entails a three-stage procedure (**Figure 1B**). In Stage 1, passive and myogenic vessel response is calibrated using pressure-diameter myography data [5]. In Stage 2, the Gharahi et al. (2023) model [24] is used to define homeostatic morphometry and hemodynamics of the three coronary subtrees. As described in the Framework Calibration Procedure section, an iterative process between Stages 1 and 2 is performed until the vessel mechanics and homeostatic quantities (hemodynamics, morphometry and SMC tone) converge. Lastly, in Stage 3, metabolic and shear-dependent autoregulatory responses are calibrated to match a given coronary pressure-flow relationship [26]. The result of this calibration procedure is a tree model endowed with morphological, hemodynamic, structural, and autoregulatory characteristics of the coronary circulation.

### 2.2 Microstructurally-motivated vessel wall model

The microstructurally-motivated wall model, which governs the passive and active behavior for all vessels, is presented here, as well the model of SMC activation and autoregulatory stimuli.

#### 2.2.1 Constrained mixture model formulation

Vessel wall mechanics are described with a CMM formulation, considering the main load-bearing constituents: elastin, collagen, and SMCs [27]. A strain energy function *w* is used to describe the relationship between stress and strain tensors in a non-linear manner for each constituent. CMMs incorporate the microstructural properties such that wall constituents are constrained to deform together but have distinct mechanical properties and stresses [28]. Each constituent *i* has a strain energy density (*w*^*i*^), mapping deformation gradients from their stress-free configuration to the *in-vivo* configuration. The total strain energy density for a vessel is the sum of its constituents: *w* = ∑_*i*_ *w*^*i*^. The function consists of passive contributions for elastic 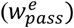 and collagen 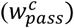, as well both passive 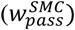 and active 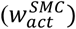 contributions for SMC. Therefore, we have:

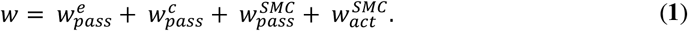

A neo-Hookean model is used for elastin (treated as an isotropic constituent with coefficient *c*_1_), and a Holzapfel’s exponential model is used for the collagen fibers (with coefficients *c*_2_ and *c*_3_) and the SMCs passive response (with coefficients *c*_4_ and *c*_5_).

Critically for this work, the active SMC response is modeled as:

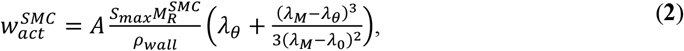

where *A* is a SMC activation level (0 < *A* < 1), *S*_*max*_ is the maximum active stress, 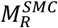 is the mass per unit area of SMCs, *ρ*_*wall*_ is the density of arterial tissue, *λ*_*θ*_ is the vessel circumferential stretch, and *λ*_0_ and *λ*_*M*_ are the zero and maximum active tension stretches, respectively. Other important material parameters in CMMs of vascular tissue are: 1) homeostatic pre-stretches at which each constituent is deposited (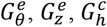, and 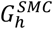), and 2) the fraction of the vessel wall occupied by each constituent (mass fractions *v*^*e*^, *v*^*c*^, and *v*^*m*^). Details on elastin, collagen, and SMC constituent orientations within the vessel wall, as well as other mathematical forms of the CMMs are summarized in **Appendix A** and available in greater detail elsewhere [23].

Using membrane theory [28], the circumferential wall tension (*T*_*θθ*_) of a thin-walled vessel is calculated from its strain energy density as:

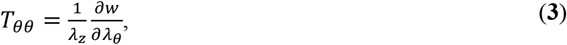

where *λ*_*z*_ is the axial stretch of the vessel. Furthermore, mechanical equilibrium is given by Laplace’s law:

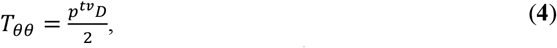

where *p*^*tv*^ is transvascular pressure and *D* is internal diameter. Note, *p*^*tv*^ is the difference between lumen pressure (*p*) and the intramyocardial pressure (*p*^*im*^): *p*^*tv*^ = *p* − *p*^*im*^. By combining Equation (**3**) and (**4**), the vessel diameter can be determined. Autoregulation is incorporated within the CMM formulation by adjusting the SMC activation *A* in response to autoregulatory stimuli.

#### 2.2.2 Modeling of autoregulatory stimulus

SMC activation is modulated between fully dilated (*A* = 0) and maximally constricted (*A* = 1). Following Carlson et al. (2008) [29], *A* is assumed to be sigmoidal and to depend on the total SMC stimulus (*s*_*total*_):

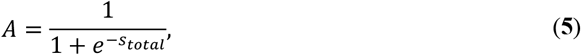

where *s*_*total*_ depends on myogenic (*s*_*myo*_), metabolic (*s*_*meta*_), and shear-dependent (*s*_*τ*_) stimuli. Furthermore, SMCs maintain a basal tone under homeostatic conditions (*s*_*h*_) [30]. Thus, *s*_*total*_ can be expressed as:

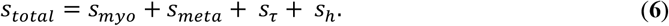

It is important to point out that under the homeostatic state, *s*_*h*_ contains contributions from each autoregulatory mechanism, even though at the homeostatic state, the three different signals (myogenic, metabolic, and shear-dependent) are zero by construction, see below. Since the relative strength of these control mechanisms depends on vessel size [2], [5], [31], [32], normalized diameter-dependent response curves for the myogenic (*ϕ*_*myo*_(*D*)), metabolic (*ϕ*_*meta*_(*D*)),), and shear-dependent (*ϕ*_*τ*_(*D*)) control mechanisms are defined below.

##### Myogenic control

The myogenic stimulus *s*_*myo*_ is assumed to be linearly dependent on wall tension *T*_*θθ*_ [29]:

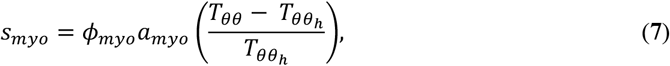

where *a*_*myo*_ is a myogenic scaling coefficient and 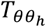 is the homeostatic wall tension. From Stage 1 model calibration, we obtain *a*_*myo*_ and *ϕ*_*myo*_ at four representative vessel sizes (*D* = 180, 95, 62, and 37 μm), as discussed in **Section 2.6.1**, using the data of Liao and Kuo [5]. To extrapolate *ϕ*_*myo*_ to other vessel diameters, we assume no myogenic response (i.e. *ϕ*_*myo*_ = 0) for the precapillary arterioles and for *D*>500 μm [26]. *ϕ*_*myo*_ is interpolated across all vessel diameters using a power-law scheme. As discussed in **Section 2.6.1**, each of the three subtrees has distinct *ϕ*_*myo*_ distributions, denoted 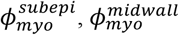, and 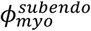.

##### Metabolic control

Changes in *MVO*_2_ is assumed to be matched by proportional changes in flow to meet the oxygen demand [33]. Based on this assumption, the metabolic stimulus *s*_*meta*_ is linearly dependent on the terminal arterioles flow rate, regardless of the underlying physiological mechanisms involved [13]:

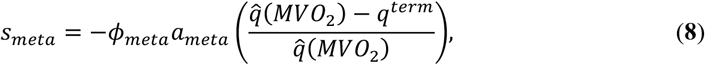

where *a*_*meta*_ is a metabolic scaling coefficient, 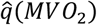 is the target flow rate determined by a baseline metabolic demand, and *q*^*term*^ is the terminal arterioles flow rate. The negative sign indicates a decrease in metabolic stimuli for a decrease in terminal flow. Furthermore, The metabolic response curve *ϕ*_*meta*_ assumes that metabolic signals originate in the capillaries and propagate upstream via endothelial gap junctions [13]:

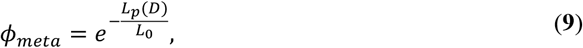

where *L*_*p*_(*D*) is the path length from the midpoint of a vessel with diameter *D* to the capillaries, and *L*_0_ is a characteristic decay length, assumed to be *L*_0_ = 0.001 m [13].

##### Shear-dependent control

An increase in wall shear stress *τ* induces relaxation of SMCs. The shear-dependent stimulus (*s*_*τ*_) can be modeled as:

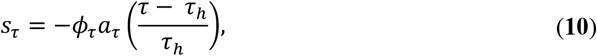

where *a*_*τ*_ is a shear-dependent scaling coefficient and *τ*_*h*_ is a homeostatic wall shear stress. The negative sign indicates a vasodilatory response from an increase in wall shear stress. Measurements of fractional dilation induced by shear stress in swine coronary arteries at four vessel sizes (*D* = 180, 95, 62, and 37 mm) from Liao and Kuo [5] were used as a proxy for the shear-dependent normalized response *ϕ*_*τ*_. Similar to myogenic control, *ϕ*_*τ*_ is extrapolated to the entire coronary tree assuming negligible shear-dependent control (i.e. *ϕ*_*τ*_ = 0) for the precapillary arterioles and for vessel dimeters ≥ 730 μm [13]. A power-law scheme is used to interpolate *ϕ*_*τ*_ across all vessel diameters.

##### Homeostatic stimulus

Note that at the homeostatic state *A* = *A*_*h*_ and *s*_*total*_ = *s*_*h*_. The homeostatic SMC stimulus *s*_*h*_ and *A*_*h*_ are related by:

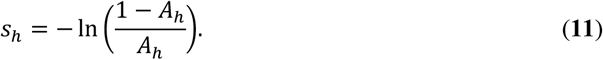

Two examples of normalized diameter-dependent response curves are presented in **Figure 2**. Once the myogenic responses are calibrated for vessels within the Liao and Kuo [5] dataset, (180 > D > 37 μm, see shaded region), two different assumptions for myogenic responses outside this range are considered. In Panel **A** (Low *ϕ*_*myo*_), the myogenic responses are assumed to drop to zero, whereas in Panel **B** (High *ϕ*_*myo*_), the myogenic responses are assumed to remain constant outside the range. Note, that the High *ϕ*_*myo*_ profile is not intended to be physiologically realistic, but is used to assess how different response curves may impact modeling results. Results are presented using the Low *ϕ*_*myo*_ profile, unless stated otherwise.

**Figure 2:**
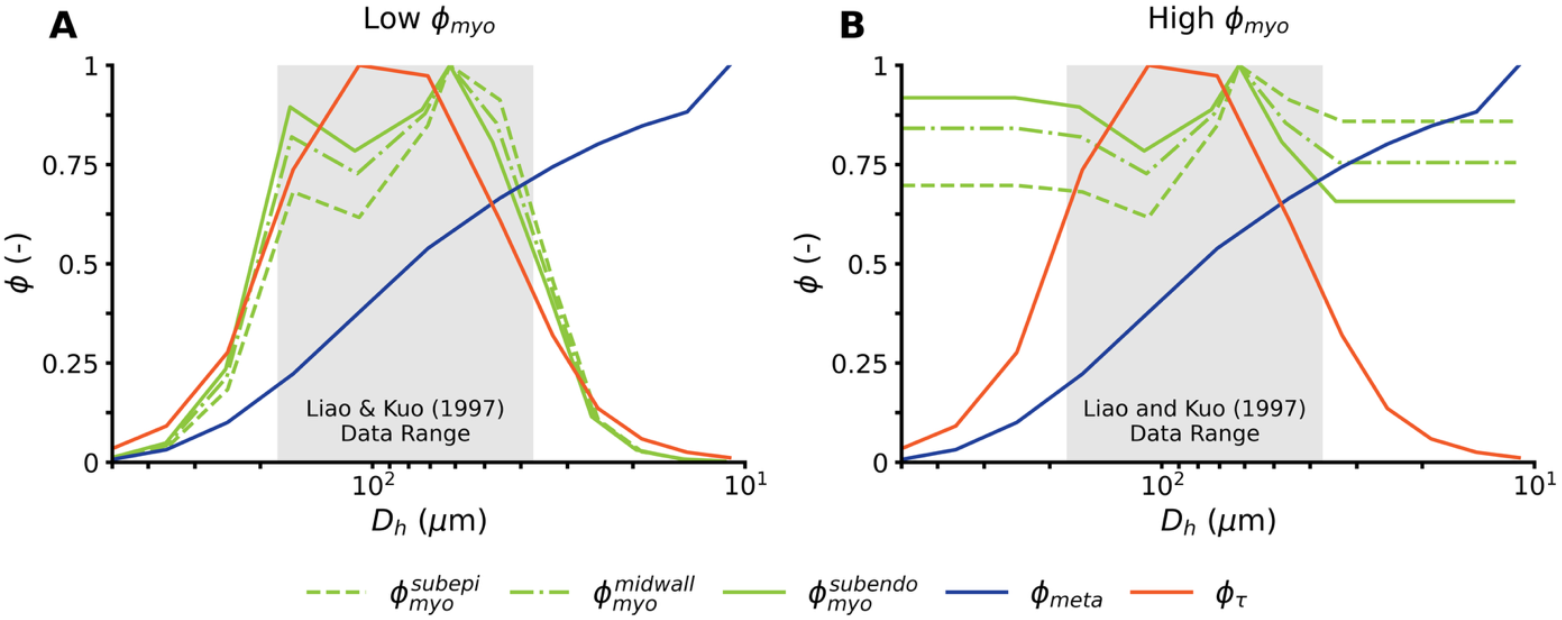
(**A**) Low *ϕ*_*myo*_ and (**B**) High *ϕ*_*myo*_ profiles for myogenic response curves. Metabolic (*ϕ*_*meta*_), and shear-dependent (*ϕ*_*τ*_) response curves are identical in both examples. The gray box indicates vessel diameters in the Liao and Kuo dataset [5].

### 2.3 A model for intramyocardial pressures

Intramyocardial pressure (*p*^*im*^) depends on a shortening-induced pressure (*SIP*) and a cavity-induced extracellular pressure (*CEP*) [34]. *SIP* accounts for myofibril contraction during myocyte activation and is assumed to remain constant across the myocardium. *CEP* is the pressure transmitted from left ventricle (*p*^*LV*^), and is greater in the endocardium than in the epicardium [35]. The distribution of *p*^*im*^ along each subtree *j* (subepi, midwall, subendo) is modeled as [15]:

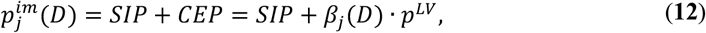

where *β*_*j*_(*D*) is a normalized myocardial depth distribution, with *β*_*j*_ = 1 at the endocardial surface and *β*_*j*_ = 0 at the epicardium surface. *SIP* and *p*^*LV*^ waveforms are taken from Namami et al. [13] (**Figure 3A**). We assign *β*_*j*_(*D*) distributions that anatomically represent coronary trees penetrating the myocardium from the epicardial surface, and approaching capillary beds located at normalized myocardial depths *β*_*cap, j*_ of 1/6, 1/2, and 5/6 for the subepi, midwall, and subendo layers, respectively:

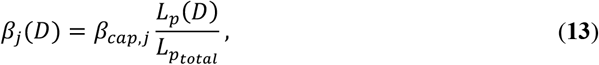

where *L*_*p*_(*D*) is the path length between the midpoint of a vessel with diameter *D* to the capillaries and 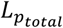 is the total subtree path length. The pulsatile *p*^*im*^ at the capillary depth *β*_*j*_(*D*) = *β*_*cap, j*_ is shown in **Figure 3A** for each coronary subtree, while the distribution of mean intramyocardial pressure 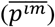 across subtree generations is presented in **Figure 3B**.

**Figure 3:**
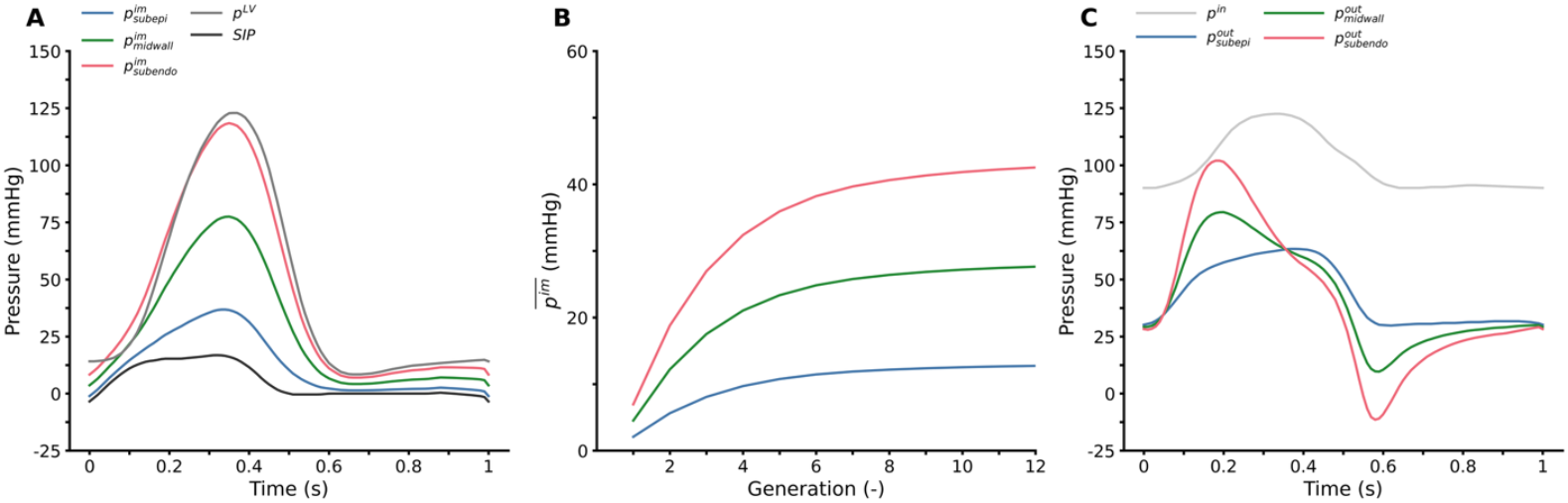
Imposed pressure waveforms on the coronary tree. (**A**) Pulsatile intramyocardial pressures (*p*^*im*^) at a normalized myocardial depth of *β*_*j*_ (*D*) = *β*_*cap, j*_ for each coronary subtree, calculated from left ventricular (*p*^*LV*^) and shortening-induced pressure (*SIP*) via **Equation (12)**. (**B**) Distribution of mean intramyocardial pressure 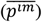 across vessel generations for each coronary subtree, based on the normalized myocardial depth distribution from **Equation (13)**. (**C**) Inlet and outlet lumen boundary pressure waveforms from Namami et al. [13].

### 2.4 Hemodynamic Modeling

Vessel fluid-solid interaction is modeled using Womersley’s solution for oscillatory flow in deformable tubes [36]. This approach solves for flow and pressure under linearized assumptions, as described in detail elsewhere [24], [37]. In this work, Womersley’s solution is extended to account for a time- and spatially-varying *p*^*im*^. Furthermore, a system of equations is constructed to solve hemodynamics throughout the coronary tree.

#### 2.4.1 Extension of Womersley’s solution for time-varying external pressure

Intramyocardial pressure is assumed to be a known periodic quantity, uniform along a single vessel of length *L* (i.e. 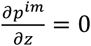). In the frequency domain, flows (*q*) and transvascular pressures (*p*^*tv*^) are related through an admittance matrix (***Y***) [38]:

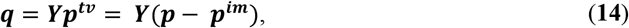

where ***q*** = [*q*_1_ *q*_2_]^*T*^ and 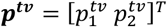, and subscripts 1 and 2 designate the vessel inlet (*z* = 0) and outlet (*z* = *L*), respectively. Details on constructing ***Y*** are presented in **Appendix B**. For non-zero frequencies (*ω* ≠ 0), ***Y*** is:

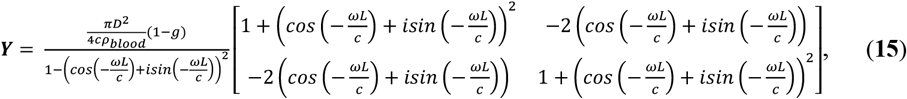

where *c* is the pulse wave velocity, *ρ*_*blood*_ is the blood density, 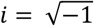, and *g* = 2*J*_1_(∧)/∧ *J*_0_(∧). *J*_0_ and *J*_1_ are Bessel functions of the first kind and 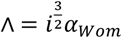, where 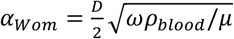 is the Womersley number and *μ* the blood viscosity. Since viscosity is dependent on vessel diameter and blood hematocrit, we utilize the *in-vivo* viscosity law given by [39] and formulated in **Appendix C**. For the steady component of Womersley’s solution (i.e. *ω* = 0), ***Y*** is [40]:

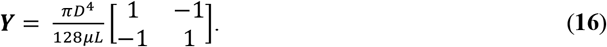

Thus, Equations (**14**)-(**16**) calculate flow and pressure and the ends of a deformable vessel.

#### 2.4.2 Womersley’s solution for an arterial network

At any bifurcation, conservation of flow and continuity of pressure states:

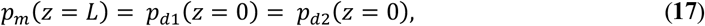

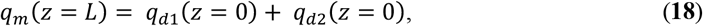

where the *m* and *d*1, *d*2 denote mother and two daughter branches, respectively. By applying Equation (**14**) to each vessel and Equations (**17**) and (**18**) at each bifurcation, we construct a linear system of equations: ***Cx*** = ***b***, where ***C*** is a coefficient matrix, ***x*** the vector of unknown flows and pressures, and ***b*** a vector containing the prescribed boundary conditions and *p*^*im*^ waveforms. In this paper, pulsatile hemodynamics are only considered at the homeostatic state, which uses the inlet and outlet pressure boundary conditions taken from Namani et al. [13] (**Figure 3C)**.

### 2.5 Framework algorithm: How to calculate an auto-regulated state

Given a fully calibrated model (passive and myogenic responses – Stage 1; homeostatic geometry and hemodynamics – Stage 2; and baseline metabolic and shear dependent responses – Stage 3), we can predict autoregulated states following a certain trigger to the system. The algorithm to predict these auto-regulated states is shown in **Figure 4**. Starting with the homeostatic tree morphometry, the admittance ***Y*** for the steady component of Womersley’s solution (i.e. *ω* = 0) is computed for all vessels (Equation (**16**)). Then, accounting for the trigger (e.g., change in perfusion pressure (*p*^*in*^)), the steady state hemodynamics are calculated throughout the coronary tree. The updated hemodynamics alter the autoregulatory stimuli (*s*_*myo*_, *s*_*meta*_, *s*_*τ*_), and consequently SMC activation *A*. Next, vessel diameters *D* throughout the tree are calculated using the updated SMC activation (Equation (**3**) and (**4**)), resulting in a new tree morphometry. The process of updating tree hemodynamics and vessel diameters is iteratively performed until convergence, determined when *p, q*, and *D* change less than a small number *ε* between two successive iterations. This algorithm outputs the autoregulated hemodynamics, tree morphometry, and autoregulatory stimuli.

**Figure 4:**
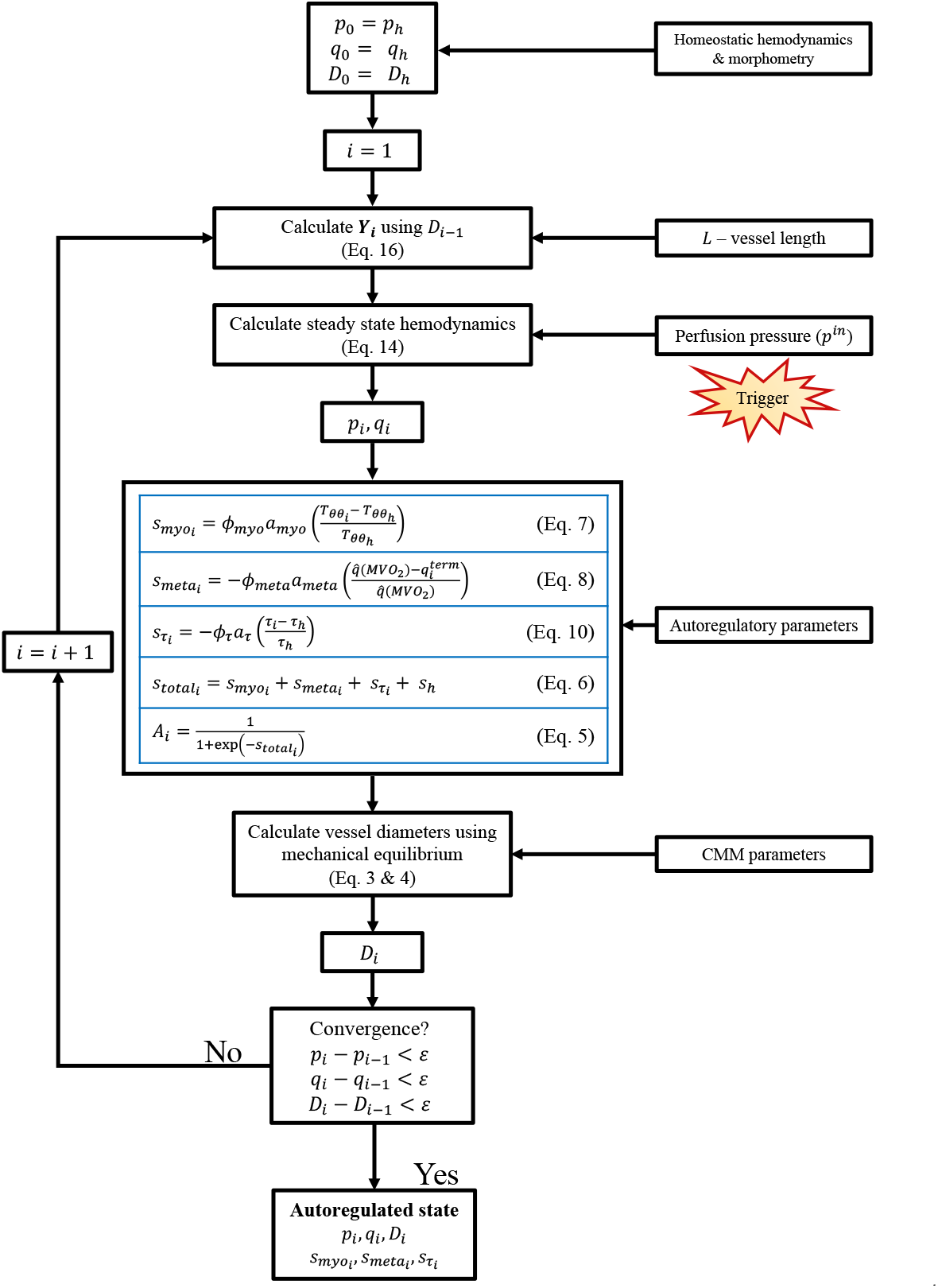
Iterative procedure to solve for the steady state coronary hemodynamics for a given perfusion pressure *p*^*in*^.

For calculating steady state hemodynamics at an autoregulated state, a constant capillary resistance *R*_*capillary*_ boundary condition, plus a 20 mmHg terminal coronary venous pressure [41], is applied to the outlets. The capillary resistance is computed as: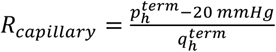, where 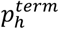 and 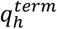 are the terminal arteriole homeostatic pressure and flow, respectively, obtained in Stage 2 model calibration (see Section 2.6.2 Stage 2 calibration: Homeostatic coronary trees).

### 2.6 Framework calibration procedure

#### 2.6.1 Stage 1 calibration: Passive and myogenic response

Myography measurements from Liao and Kuo are used to determine the passive and myogenic response of vessels 180 μm (small arteries), 95 μm (large arterioles), 62 μm (intermediate arterioles), and 37 μm (small arterioles) in diameter [5]. Identical myogenic responses are assumed for trees in the different myocardial depths (subepi, midwall, and subendo). This assumption is made given conflicting evidence regarding transmural differences in myogenic strength across the myocardium [26], [42], [43] within the experimental pressure range (15-100 mmHg).

To describe the passive response of the vessel, the active SMC contribution is removed from the strain energy function (i.e. 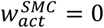), see **Equation (1)**. Conversely, the purely myogenic response is modeled by removing the metabolic and shear-dependent stimuli in **Equation (6)**, i.e., *s*_*meta*_ = *s*_*τ*_ = 0, and considering a SMC stimulus 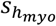 which neglects the metabolic and shear-dependent contributions embedded in *s*_*h*_. Thus, the total myogenic stimulus is defined as:

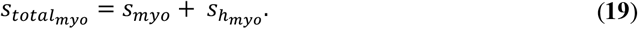

Passive and myogenic response calibration is performed under the assumption that mechanical properties of constituents (material stiffnesses *c*_1_−*c*_5_ and active SMC parameters *S*_*max*_, *λ*_*M*_, *λ*_0_) are constant across the vascular tree, but mass fractions, homeostatic deposition stretch of different constituents (pre-stretches), and myogenic strength depend on vessel size and myocardial depth [24], [26]. Overall, identifying CMM parameters of 12 representative vessels (four vessels in each myocardial depth) requires estimating 80 parameters. Several physiological assumptions were made to avoid unrealistic values of model parameters. Constituent pre-stretches <1 are not allowed in the model. Collagen fibers are assumed to have a relatively low pre-stretch 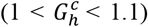, whereas elastin and SMC are allowed to have larger pre-stretches (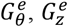 and 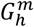, respectively) [44]. These assumptions reflect the high turnover of collagen (leading to low pre-stretch) compared to the much lower turnover of elastin and SMC. Furthermore, collagen fiber angles were prescribed to follow axial, circumferential, and diagonal directions (e.g. 0°, 90° and 45°, respectively). Additionally, a homeostatic SMC activation (*A*_*h*_) of 0.5 is targeted [29], assuming an equal vasodilatory and vasoconstrictive capacity at the homeostatic state.

The error function for passive and myogenic calibration for a vessel is given by:

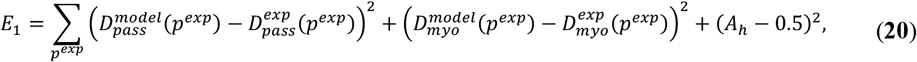

where, *p*^*exp*^ is the set of experimental pressures tested by Liao and Kuo [5], and the different *D* terms refer to vessel diameters under different conditions: *pass* and *myo* indicate the purely passive and myogenic responses, respectively, while *model* and *exp* indicate model and experimental diameters, respectively. Liao and Kuo do not report a homeostatic pressure (*p*_*h*_) or vessel thickness (*H*_*h*_), both necessary for CMM parameter calibration [5]. Therefore, we initially estimated them from literature [20] and iteratively updated them using the homeostatic trees determined in Stage 2 calibration. A Nelder-Mead algorithm was used to minimize the error function in equation (**20**). Details are provided in **Appendix D**.

#### 2.6.2 Stage 2 calibration: Homeostatic coronary trees

The homeostatic morphometric (e.g., diameters), structural (e.g., thickness), and hemodynamic (e.g., wall shear stress) properties of the subtrees are determined using the homeostatic optimization model presented by Gharahi et al. [24]. The homeostatic values of two specific vessel properties, vessel diameter *D*_*h*_ and SMC activation *A*_*h*_ are obtained using cycle-averaged hemodynamic quantities (i.e., the steady component in Womersley’s solution **Section 2.4**).

Each subtree is assumed to be symmetrically bifurcating over 12 generations, with the first generation having a homeostatic diameter *D*_*h*_ = 500 μm (**Table 1**). Constant inlet pressure and constant outflow boundary conditions are prescribed for each subtree (**Table 1**). Vessel lengths were determined as a function of vessel diameters using the morphometric data from Guo and Kassab [20].

**Table 1:** Parameters for coronary homeostatic optimization.

| Parameter | Description | Unit | Value | Source |
| --- | --- | --- | --- | --- |
| $N$ | Number of generations | - | 12 | - |
| $D_0$ | Diameter of first generation | $\mu\text{m}$ | 500 | - |
| $p_h^{\text{in}}$ | Homeostatic coronary inlet pressure | mmHg | 100 | [45] |
| $q_{\text{subepi}}^{\text{term}}$ | Subepi homeostatic terminal flow | $\text{mm}^3/\text{s}$ | 0.001375 | [13], [15] |
| $q_{\text{midwall}}^{\text{term}}$ | Midwall homeostatic terminal flow | $\text{mm}^3/\text{s}$ | 0.0015 | [13] |
| $q_{\text{subendo}}^{\text{term}}$ | Subendo homeostatic terminal flow | $\text{mm}^3/\text{s}$ | 0.00165 | [13], [15] |
| $p_{\text{subepi}}^{\text{im}}$ | Subepi mean intramyocardial pressure | mmHg | 12.7 | [13] |
| $p_{\text{midwall}}^{\text{im}}$ | Midwall mean intramyocardial pressure | mmHg | 27.6 | [13] |
| $p_{\text{subendo}}^{\text{im}}$ | Subendo mean intramyocardial pressure | mmHg | 42.5 | [13] |
| $\rho_{\text{blood}}$ | Blood density | $\text{kg}/\text{m}^3$ | 1060 | - |

CMM parameters determined in Stage 1 calibration are assigned to individual vessels according to their homeostatic diameter *D*_*h*_ (Small Arteries: *D*_*h*_>140 μm, Large Arterioles: 140≥*D*_*h*_>80 μm, Intermediate Arterioles: 80≥*D*_*h*_>50 μm, Small Arterioles: 50≥*D*_*h*_ μm) and myocardial depth. However, grouping vessels into four classes is crude, since considerable hemodynamic and morphological variability may exist within one vessel class. We have observed that when CMM parameters (specifically SMC tone) are kept constant across all classes, homeostatic optimization does not produce the experimentally measured thickness-diameter relationship. Therefore, we calibrate the homeostatic SMC activation parameter *A*_*h*_ (**Equation (11)**) such that each vessel attains its target thickness *H*_*h*_. **Appendix A** outlines the mathematical relationship between *A*_*h*_ and *H*_*h*_, following mechanical equilibrium principles stated in Equations (**3**) and (**4**)).

Using measured thickness-diameter relationships for subepicardial vessels [20], the following strategy is used to prescribe the target thickness *H*_*h*_ for individual vessels. First, *A*_*h*_ values in the subepi tree were adjusted, and the homeostatic hoop stress (*σ*_*h*_) was obtained through: 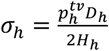. Then, *A*_*h*_ in the midwall and subendo layers was adjusted to match the subepi hoop stress distribution, based on the hypothesis proposed by Choy and Kassab [9] that hoop stress remains uniform across the myocardial wall for a given vessel size. This hypothesis is supported by experimental measurements showing transmural differences in coronary vessel wall thickness (e.g., thicker walls are found in the subepi region for vessels of similar diameters).

A key morphometric output of the homeostatic optimization is the bifurcation exponent *ξ*, which describes the diameter relationship between mother (*D*_*m*_) and daughter (*D*_*d*1_ and *D*_*d*2_) vessels at each bifurcation as: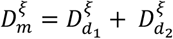. Kaimovitz et al. [46] reported *ξ* throughout the swine coronary circulation and found considerable heterogeneity, likely reflecting the size-dependent functional roles of coronary vessels [2]. In this work, we add a penalty term to the cost function used by Gharahi et al. [24] to incorporate the *ξ* measurements by Kaimovitz et al. [46]. Details on the cost function formulation for homeostatic optimization are provided in **Appendix E**.

#### 2.6.3 Stage 3 calibration: Metabolic and shear-dependent control

Calibration Stage 3 determines the metabolic *a*_*meta*_ (**Equation (8)**) and shear-dependent *a*_*τ*_ (**Equation (10)**) scaling coefficients using pressure-flow autoregulation data in swine [26], while keeping the myogenic scaling coefficient *a*_*myo*_ fixed. Nelder-Mead optimization is performed to determine *a*_*τ*_ and *a*_*meta*_ such that the following error function is minimized:

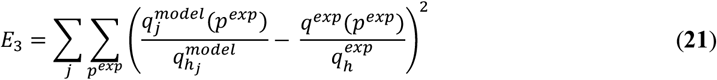

where *j* ∈ {subendo, midwall, subepi}, *p*^*exp*^ is the set of experimental pressures tested [26], *q*_*j*_ is the total flow to each coronary subtree *j*, and 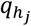 is the homeostatic flow going to coronary subtree *j*. The superscripts *model* and *exp* indicate the model and experimental flows, respectively. For a set of *a*_*meta*_ and *a*_*τ*_, the regulated coronary flow 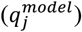 is calculated using the iterative procedure discussed in Section 2.5 and illustrated in **Figure 4**.

### 2.7 Model sensitivity

A sensitivity analysis is performed to assess model parameter sensitivity using the approach of Pradhan et al. [12]. A sensitivity index *X* was defined as the percentage change in the error function *E*_3_ (**Equation (21)**) resulting from a 10% change in individual parameter values:

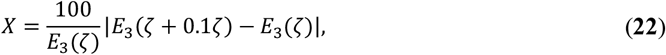

where *ζ* is the parameter value and *E*_3_(*ζ*) is its associated error. Note, this arbitrary choice of error function is chosen to assess the impact of a given parameter on global functional flow-pressure relationship and does not consider the error functions used in Stage 1 or Stage 2 of model calibration. This choice in error function emphasizes the autoregulation component of the framework over the myogenic and homeostatic components.

## 3. RESULTS

In this section, we present results from our model calibration strategy and demonstrate its application through several examples.

- <u>Model calibration and homeostatic predictions</u>: First, in **Section 3.1**, we review the three calibration stages and compare the passive, active, homeostatic, and autoregulatory properties of our coronary tree to literature data. In **Section 3.2**, we present homeostatic pulsatile hemodynamic simulations at three myocardial depths and in a small artery and small arteriole.
- <u>Autoregulatory responses to changes in perfusion pressure</u>: **Section 3.3** showcases the autoregulatory responses to changes in coronary perfusion pressure, exploring the relative magnitudes of autoregulatory stimuli. In addition, we compare the transmural autoregulatory behavior and vessel diameter changes with experimental data not used in model calibration.
- <u>Sensitivity studies</u>: In **Section 3.4**, relative contributions of the three autoregulatory mechanisms in coronary pressure-flow autoregulation are examined. Finally, in **Section 3.5**, we assess how changes in passive and active vessel mechanics affect coronary pressure-flow autoregulation.

### 3.1 Framework calibration

#### 3.1.1 Stage 1 calibration: Passive and myogenic response

Our calibrated vessel wall model effectively captures experimentally measured passive and myogenic pressure-diameter relationships across all vessel sizes and myocardial depths [5] (**Figure 5A-D**). Calibrated model parameters are given in **Table 2, Table 3**, and **Figure 5E**. The passive responses (dashed lines) are independent of myocardial depth, whereas the myogenic responses (continuous lines) exhibit clear depth-dependent differences at *p*^*tv*^>100 mmHg (**Figure 5A-D**). For each vessel class, the subendo layer has the greatest mass fraction of SMCs (**Figure 5E**). Notably, large and intermediate arterioles have the largest SMC mass fractions, consistent with having the most pronounced myogenic response [5].

**Table 2:** Constrained mixture model parameters that are constant across vessel sizes.

| Parameter | Description | Unit | Value | Sensitivity (%) | Source |
| --- | --- | --- | --- | --- | --- |
| **Global** |  |  |  |  |  |
| $c_1$ | Elastin material parameter | Pa/kg | 94.5 | 1.42 | Calibrated (Stage 1) |
| $c_2$ | Collagen material parameter | Pa/kg | 308.6 | 0.21 | Calibrated (Stage 1) |
| $c_3$ | Collagen dimensionless material parameter | - | 3.9 | 0.64 | Calibrated (Stage 1) |
| $c_4$ | SMCs material parameter | Pa/kg | 57.2 | 0.30 | Calibrated (Stage 1) |
| $c_5$ | SMCs dimensionless material parameter | - | 0.59 | 0.65 | Calibrated (Stage 1) |
| $S_{max}$ | SMC maximum stress | kPa | 250 | 9.52 | [23], [47] |
| $\lambda_0$ | Zero active tension stretch | - | 0.4 | 0.11 | [23], [47] |
| $\lambda_M$ | Maximum active tension stretch | - | 1.1 | 3.23 | [23], [47] |
| $\rho_{wall}$ | Vessel density | kg/m<sup>3</sup> | 1060 | - | - |
| $L_0$ | Metabolic control characteristic decay length | m | 0.001 | - | [13] |
| **Subepi** |  |  |  |  |  |
| $a_{myo}$ | Myogenic scaling coefficient | - | 6.32 | 0.01 | Calibrated (Stage 1) |
| $a_{meta}$ | Metabolic scaling coefficient | - | 2.82 | 0.87 | Calibrated (Stage 3) |
| $a_\tau$ | Shear-dependent scaling coefficient | - | 1.58 | 0.68 | Calibrated (Stage 3) |
| **Midwall** |  |  |  |  |  |
| $a_{myo}$ | Myogenic scaling coefficient | - | 5.43 | 0.67 | Calibrated (Stage 1) |
| $a_{meta}$ | Metabolic scaling coefficient | - | 2.82 | 0.87 | Calibrated (Stage 3) |
| $a_\tau$ | Shear-dependent scaling coefficient | - | 1.58 | 0.68 | Calibrated (Stage 3) |
| **Subendo** |  |  |  |  |  |
| $a_{myo}$ | Myogenic scaling coefficient | - | 4.37 | 0.09 | Calibrated (Stage 1) |
| $a_{meta}$ | Metabolic scaling coefficient | - | 2.82 | 0.87 | Calibrated (Stage 3) |
| $a_\tau$ | Shear-dependent scaling coefficient | - | 1.58 | 0.68 | Calibrated (Stage 3) |

**Table 3:**
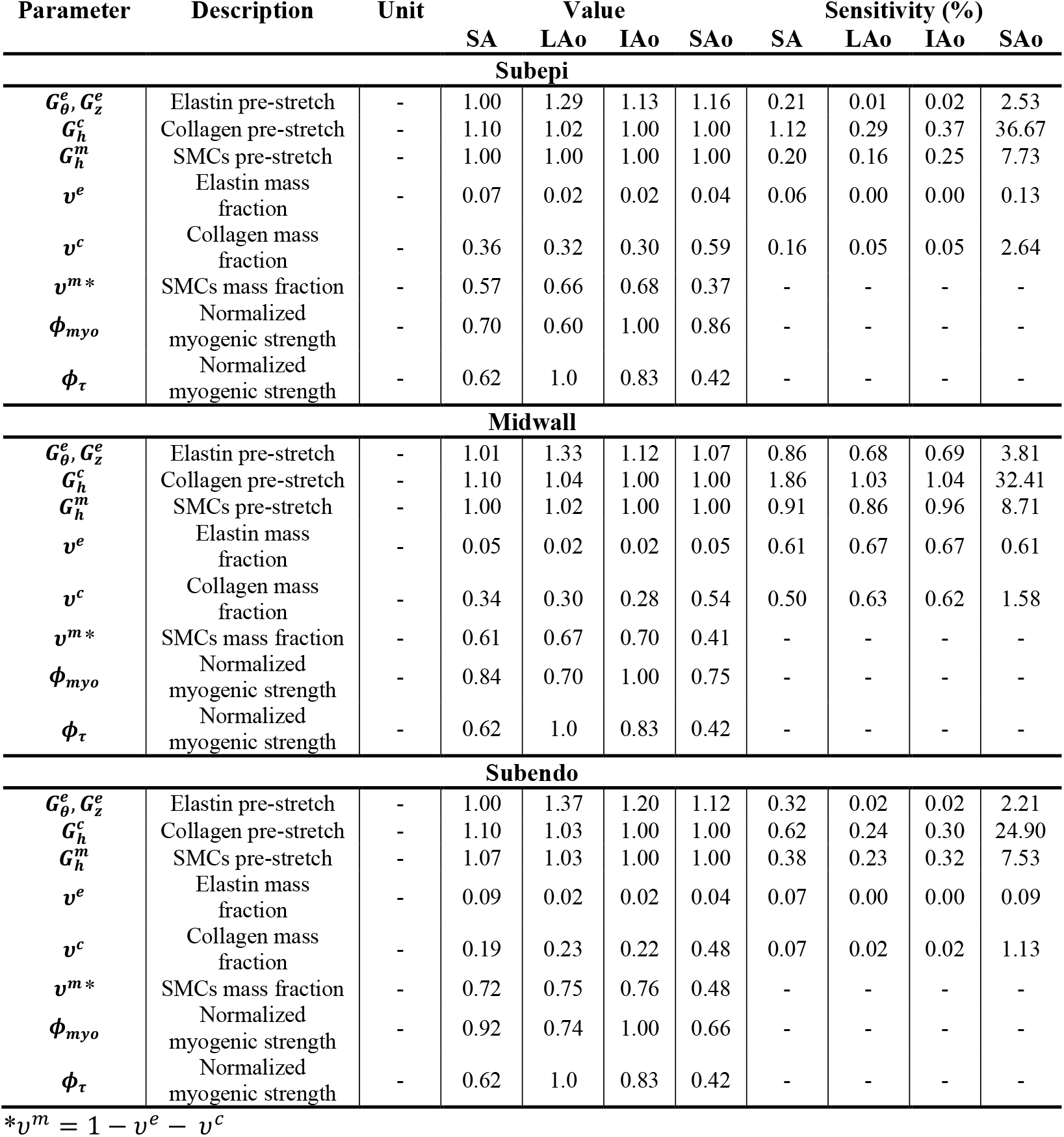
Constrained mixture model parameters that are variable across vessel sizes. small artery: SA, large arteriole: Lao, intermediate arteriole: IAo, and small arteriole: SAo.

**Figure 5:**
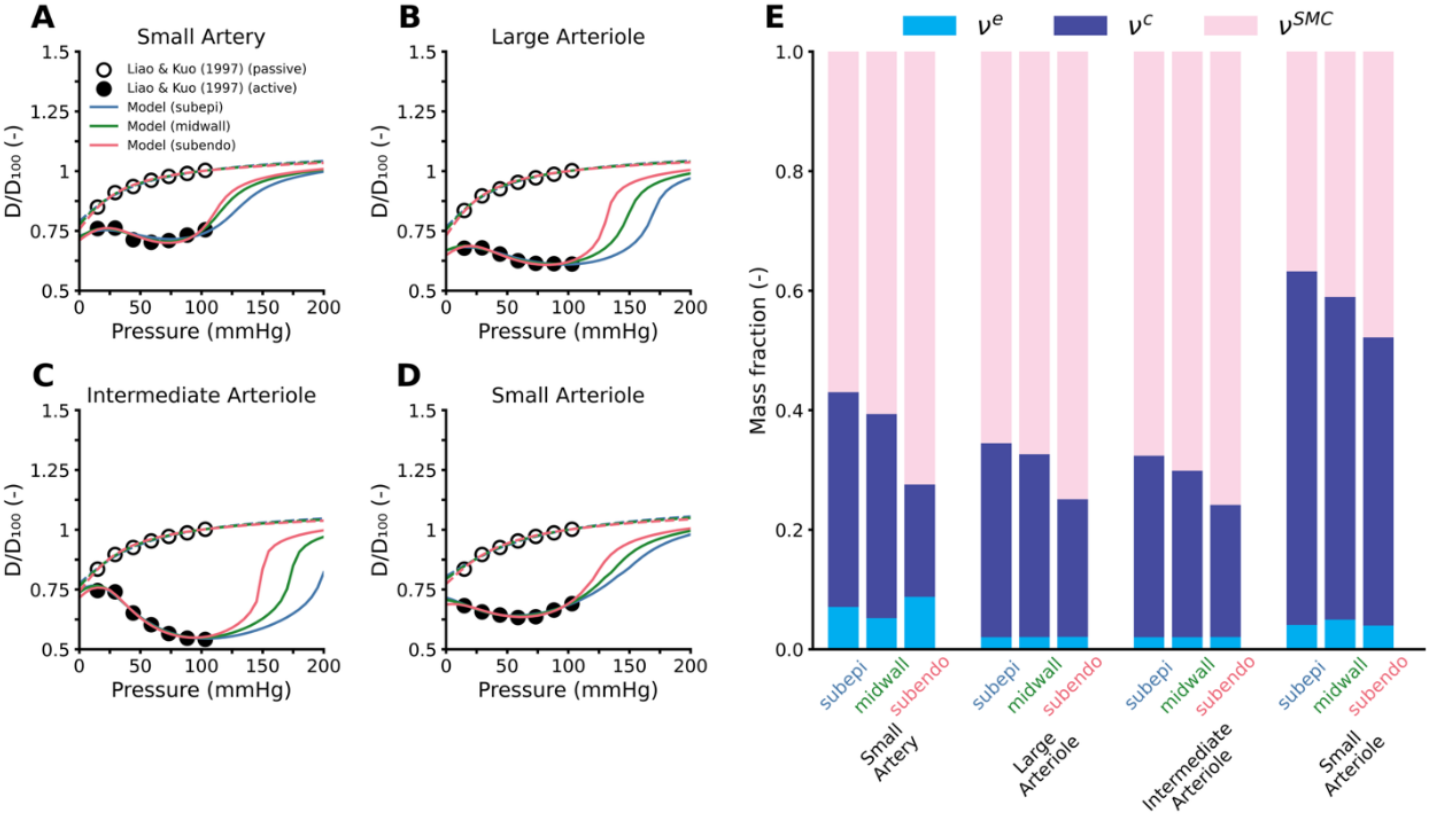
Calibrated passive and myogenic responses across four sizes of coronary vessels (**A**: Small Artery, **B**: Large Arteriole, **C**: Intermediate Arteriole, **D**: Small Arteriole), compared to experimental data from [5]. The y-axis shows diameter normalized by the passive diameter at 100 mmHg (D/D_100_). Estimated mass fractions of elastin (*v*^*e*^), collagen (*v*^*c*^), and smooth muscle cells (*v*^*SMC*^) vary across vessel sizes and myocardial depths (**E**).

#### 3.1.2 Stage 2 calibration: Homeostatic morphometry, structure, and hemodynamics

Homeostatic optimization results are summarized in **Figure 6**. Vessel diameters across generations show minimal dependence on myocardial depth (**Figure 6A**). Consequently, diameter bifurcation exponent *ξ* is also not depth dependent (**Figure 6B**) and is qualitatively aligned with measurements by Kaimovitz et al. [46], though our model predicts larger *ξ* exponents for D>65 μm. The vessel length-diameter relationship matches that presented by Guo and Kassab [20] and does not differ between subtrees (**Figure 6C**).

**Figure 6:**
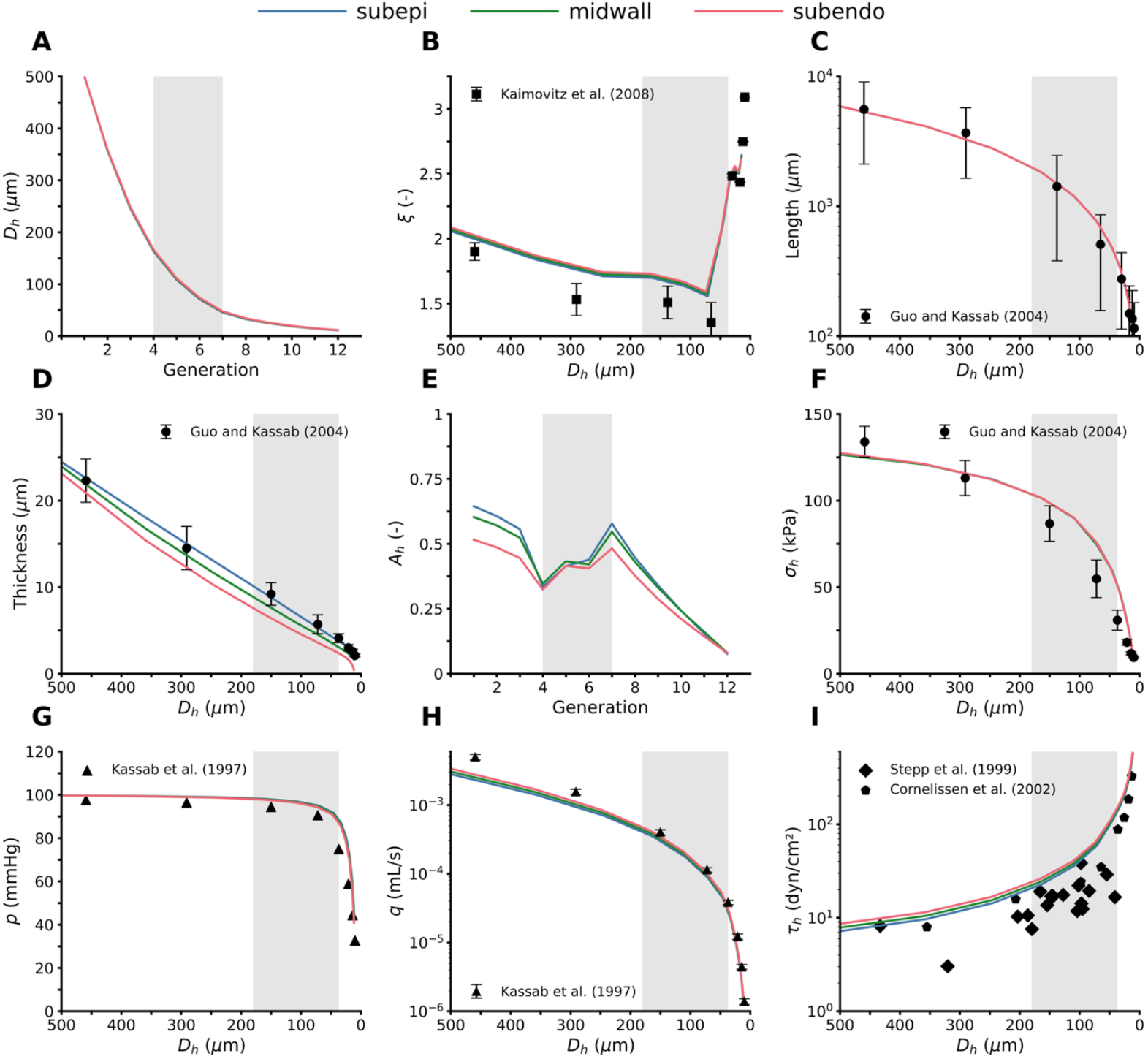
Comparison of coronary trees generated through the homeostatic optimization framework (Stage 2 calibration). The three tree morphologies have little difference in their diameter (**A**), diameter exponent (**B**), and vessel length (**C**) distributions. However, there is a layer dependence on vessel thickness (**D**). The homeostatic SMC activation (**E**) was found to vary between the three layers when targeting equivalent hoop stress distributions (**F**). Mean pressure (**G**), flow (**H**), and shear stress (**I**) vary slightly because of different terminal flows being prescribed for the three tree layers. The gray box indicates vessel diameters in the Liao and Kuo dataset [5]. Literature data are presented as mean ± SD.

Subepi tree thickness (blue line in **Figure 6D**) was calibrated using experimental data by Guo and Kassab [20] through calibration of the SMC homeostatic activation parameter *A*_*h*_ (**Figure 6E**). This produced a hoop stress distribution, which was then assumed to be identical across all three subtrees (**Figure 6F**). Using this assumption, distributions of thickness and SMC homeostatic activation *A*_*h*_ could be determined for the midwall and subendo trees (green and red lines in **Figure 6D** and **Figure 6E**, respectively). Subepi vessels are thicker than subendo vessels, consistent with anatomical measurements [9]. *A*_*h*_ was highest in large vessels and lowest in small vessels (**Figure 6E**). Of note, the vessel sizes in the Liao and Kuo (1997) data set used in Stage 1 calibration span generations 4-7 [5], where noticeable changes in *A*_*h*_ occur. Additionally, *A*_*h*_ was lowest in the subendo layer, while the subepi and midwall layers exhibited similar activation levels. The hoop stress distributions across all three coronary layers demonstrate fair agreement with Guo and Kassab [20] (**Figure 6F**).

Pressure and flow distributions are presented in **Figure 6G** and **Figure 6H**, respectively, and are comparable to the pressure and flow distributions calculated by Kassab et al. [45]. Most of the vascular resistance is in arterioles with *D*_*h*_<100 μm, as shown by the rapid decrease in pressure in **Figure 6G**. Wall shear stress increases roughly 80-fold across the coronary trees (**Figure 6I**), with values calculated by our model tending to be greater than that reported by Stepp et al. [32] but in fair agreement with the mathematical model presented by Cornelissen et al. [11].

Collectively, **Figure 6** shows that our homeostatic optimization framework produced a coronary tree that is morphologically and hemodynamically consistent with what has previously been reported in the literature.

#### 3.1.3 Stage 3 calibration: Coronary pressure-flow response

Calibrated values of metabolic *a*_*meta*_ and shear-dependent *a*_*τ*_ scaling coefficients are presented in **Table 2**. Furthermore, the autoregulated, fully dilated (*A* = 0), and maximally constricted (*A* = 1) flow-pressure relationships are presented in **Figure 7A**, with the ratio of total flow (*q*) over homeostatic flow (*q*_*h*_) in the coronary arteries plotted over varying inlet perfusion pressures *p*^*in*^.

**Figure 7:**
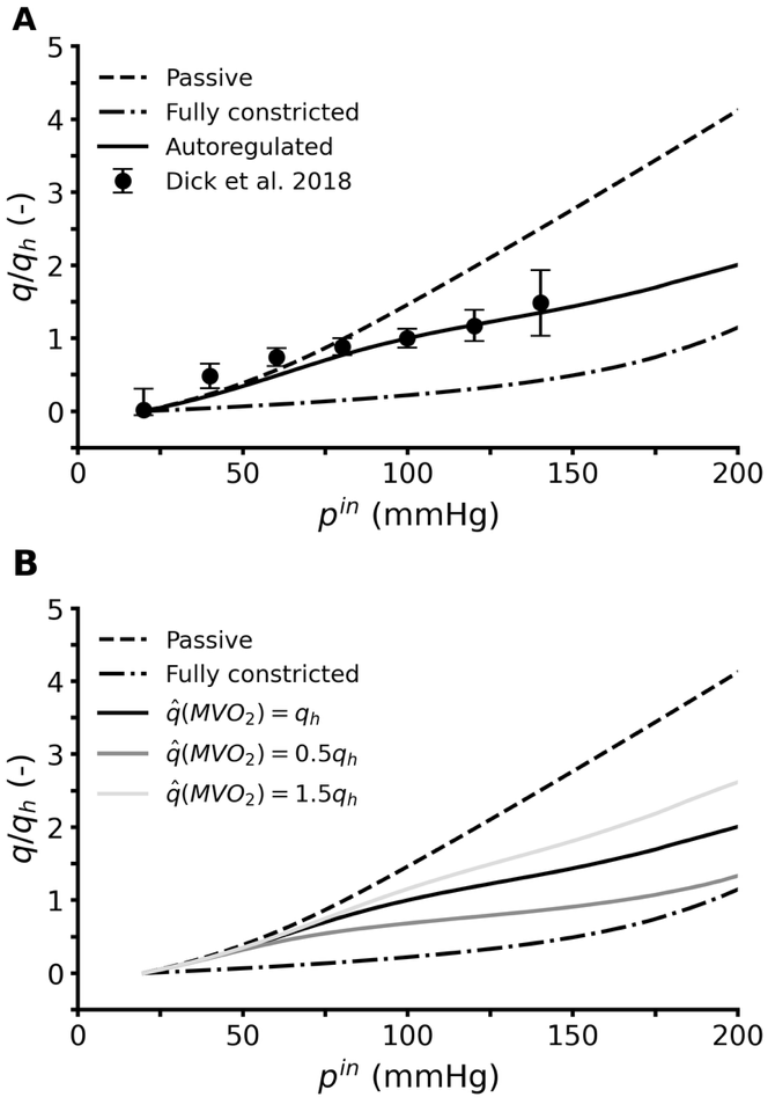
(**A**) The calibrated autoregulated pressure-flow relation for the entire tree (solid line) and is compared to experimental measurements [26], with *q*_*h*_ being the homeostatic flow rate. (**B**) A 50% increase or decrease in myocardial metabolic demand shifts the coronary pressure-flow relationship vertically up or down, respectively. Literature data are presented as mean ± SD.

At *p*^*in*^ ∈ (80 − 140) mmHg, our model fits the experimentally reported flow-pressure relationship well. However, our model underpredicts flow at *p*^*in*^ of 40 and 60 mmHg. Note, in this pressure range the predicted autoregulation curve is closely aligned to the fully dilated pressure-flow relation, indicating a full dilation of the resistive vessels. To assess how our model responds to changes in metabolic demand, we computed flow-pressure relationships in response to 50% increase 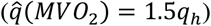 and 50% decrease 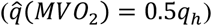 in target coronary flow (see **Equation (8)**). This alters the metabolic stimulus *s*_*meta*_, see **Figure 4**. As expected, the flow-pressure curves show upwards and downwards shifts in response to increases and decreases in metabolic demands, respectively, see **Figure 7B**.

### 3.2 Homeostatic coronary flow simulations

Coronary flow and transvascular pressure waveforms are shown for a small artery (*D*_*h*_ = 246 *μ*m, circular marker in **Figure 8A**) and small arteriole (*D*_*h*_ = 25 *μ*m, square marker in **Figure 8A**) for subepi, midwall, and subendo subtrees. Flow and pressure pulsatility is greatest in the subendo layer for both vessel sizes (**Figure 8B-E**), attributable to the intramyocardial pressure (*p*^*im*^) waveform having the greatest pulse pressure in the subendo layer (**Figure 3A**). Flow is diastolically dominant for both vessel sizes in the midwall and subendo subtrees. However, blood flow is more evenly distributed throughout the cardiac cycle in the subepi layer, with 53.3% and 47.8% of total flow occurring during systole for the small artery and small arteriole, respectively (**Figure 8B,C**). Retrograde flow occurs in all subtrees for the small artery, with the subendo layer having the most retrograde flow and the subepi layer having the least (**Figure 8B**). Conversely, the small arteriole exhibits no retrograde flow in the subepi and midwall layers, and only minimal retrograde flow in the subendo layer (**Figure 8C**). Notably, the shape of the subepi transvascular pressure waveforms differs from the midwall and subendo waveforms during systole and early diastole (**Figure 8D,E**), which ultimately impacts flow dynamics.

**Figure 8:**
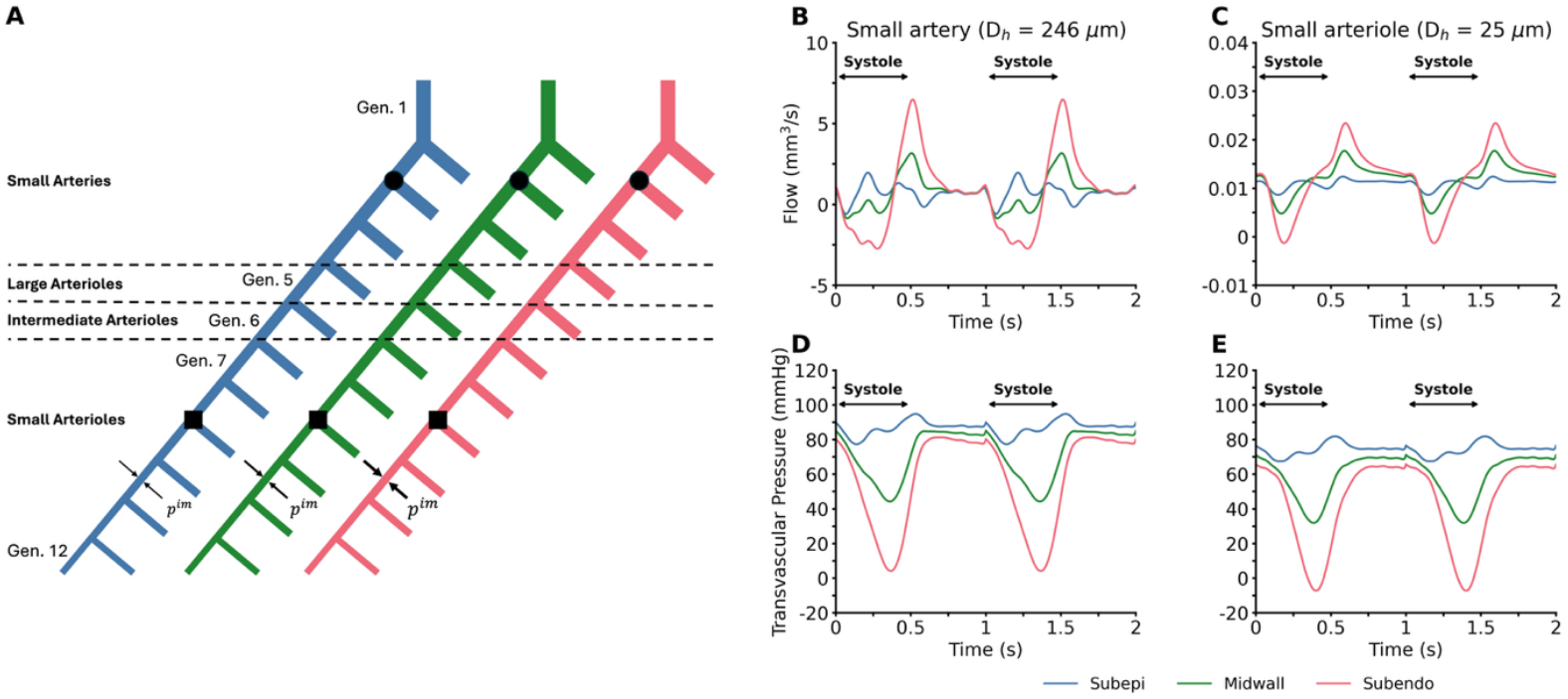
Graphical representation of the three coronary subtrees (A). The circular and square markers indicate the location on the subtrees the pulsatile flow (B-C) and transvascular pressures (D-E) are taken from, respectively.

### 3.3 Autoregulation in response to changes in perfusion pressure

#### 3.3.1 Pressure, diameter, wall shear stress, and autoregulatory stimuli

Pressure, diameter, wall shear stress, autoregulatory stimuli, and SMC activation (*A*) are presented for varying perfusion pressures in a large arteriole (*D*_*h*_ = 110 *μ*m; **Figure 9A-G**) and small arteriole (*D*_*h*_ = 25 *μ*m; **Figure 9H-N**).

**Figure 9:**
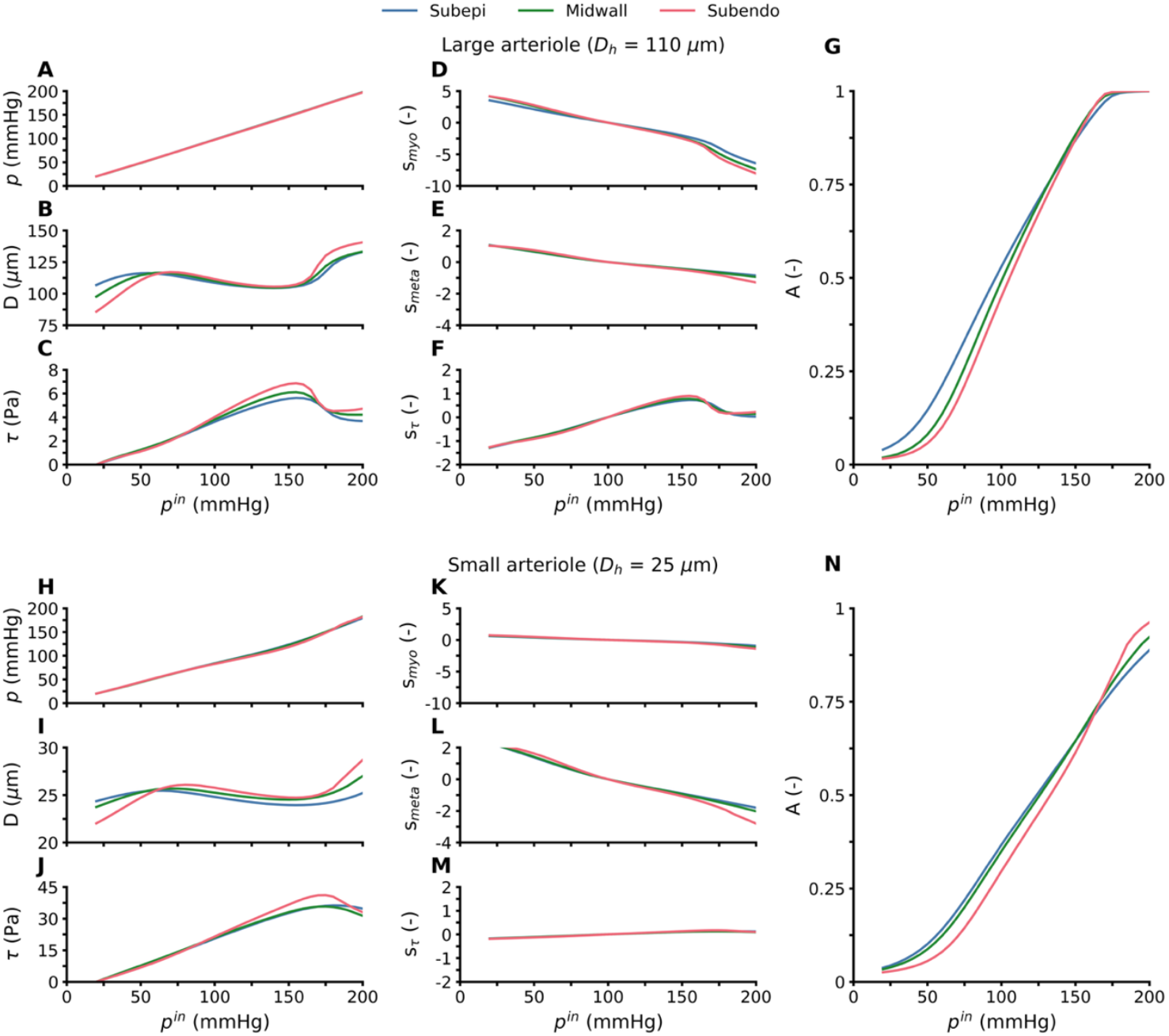
Simulated pressure, diameter, wall shear stress, autoregulatory stimuli, and SMC activation (*A*) for varying perfusion pressures in a large arteriole (*D*_*h*_ = 110 *μ*m; **Figure 9A-G**) and small arteriole (*D*_*h*_ = 25 *μ*m; **Figure 9H-N**).

Lumen pressure (*p*) increases linearly with perfusion pressure (*p*^*in*^) in the large arteriole (**Figure 9A**), but exhibits a nonlinear increase in the small arteriole (**Figure 9H**). For both vessel sizes, *p* shows minimal variation between the layers. Diameter (*D*) responses show both vessel sizes modulating their diameters over the autoregulatory pressure range, with vessel diameters decreasing as *p*^*in*^ increases from approximately 60 to 165 mmHg (**Figure 9B,I**). Outside this range, diameters increase with increasing pressure, consistent with the mechanical behavior in vessels not actively modulating tone. There are notable differences in the diameter-pressure response between the three layers, with the subendo and subepi layers actively modulating their diameters within the narrowest and widest pressure ranges, respectively, for both vessel sizes. Wall shear stress (*τ*) increases with perfusion pressure in both vessel sizes until approximately 165 mmHg before declining (**Figure 9C,J**), which is consistent with the sharp increase in diameter. The change in wall shear stress is layer-dependent; however, it is qualitatively consistent between the layers.

The large arteriole exhibits predominantly myogenic control (Figure 9D), with the myogenic stimulus greatest in the subendo layer for *p*^*in*^<100 mmHg and lowest for *p*^*in*^>100 mmHg. Metabolic and shear-dependent contributions are comparatively small and show minimal layer dependence (**Figure 9E,F**). In contrast, the small arteriole shows greater metabolic control (**Figure 9L**), with myogenic control contributing substantially less than in the large arteriole (**Figure 9K**). In small arterioles, the metabolic stimulus is greatest in the subendo layer for *p*^*in*^<100 mmHg and lowest for *p*^*in*^>100 mmHg, whereas myogenic and shear-dependent stimuli remain relatively uniform across layers (**Figure 9K,M**). The larger metabolic stimulus and smaller myogenic stimulus for the small arteriole are a consequence of the assumed normalized response curves in **Figure 2A**. The *A* versus *p*^*in*^ relationship is steepest in the subendo layer for both vessel sizes (**Figure 9G,N**), indicating vessels in this layer reach maximum dilation (A = 0) or constriction (A = 1) within a narrower pressure range. This is consistent with the diameter responses (**Figure 9B,I**), where the subendo layer exhibits the narrowest autoregulatory pressure range.

#### 3.3.2 ENDO/EPI responses

The ratio of subendo to subepi flow (ENDO/EPI) at varying perfusion pressures is shown in **Figure 10A** for low and high *ϕ*_*myo*_ normalized response curves (**Figure 2)**. From an assumed homeostatic ENDO/EPI ratio of 1.2 at *p*^*in*^=100 mmHg (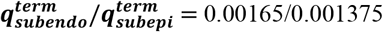, **Table 1**), the low *ϕ*_*myo*_ profile produces increased ENDO/EPI ratios for *p*^*in*^>100 mmHg. Conversely, for *p*^*in*^<100 mmHg, we see a decrease in ENDO/EPI ratio, indicative of the subendo layer being at greater risk of ischemia during hypotension. These general trends have previously been reported in canine experiments, both anesthetized [10], [48] and un-anesthetized [47]. When the high *ϕ*_*myo*_ myogenic profile is utilized, a profound difference on ENDO/EPI ratio relative to low *ϕ*_*myo*_ profile is observed for *p*^*in*^>100 mmHg, but not for *p*^*in*^<100 mmHg.

**Figure 10:**
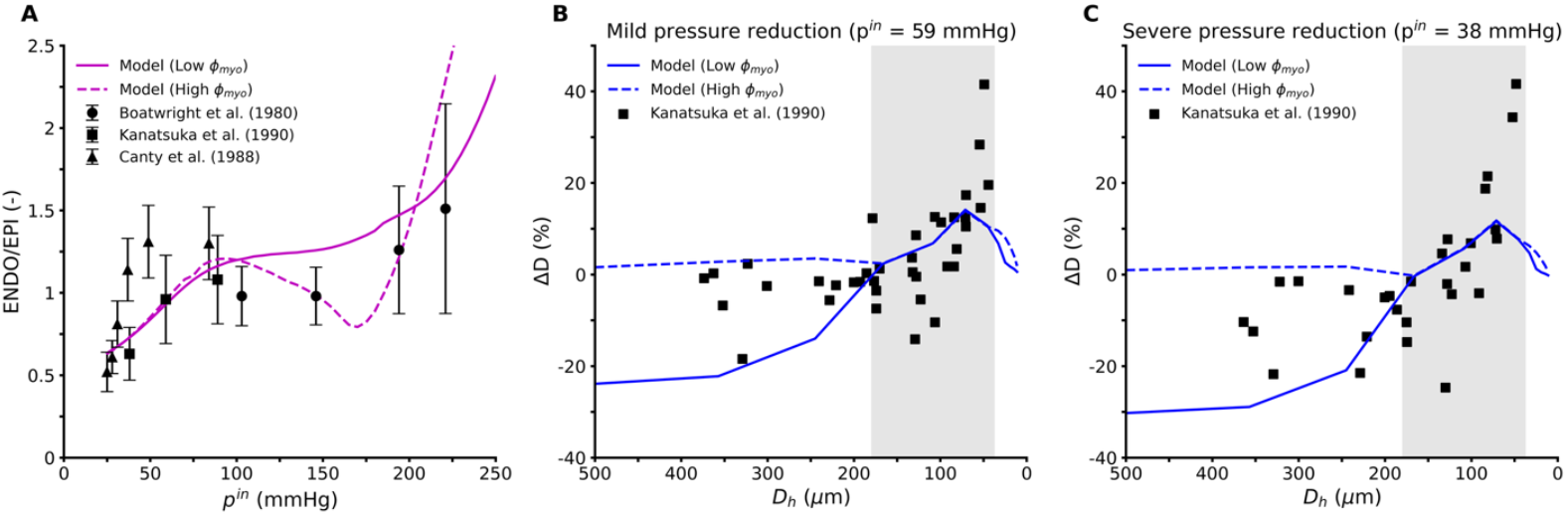
The response of our model at varying inlet perfusion pressures (*p*^*in*^). (**A**) The calculated ratio of flow going into the subepi layer vs. subendo layer (ENDO/EPI) matches the trends reported in literature. Furthermore, our models shown a dilation of arterioles in response to a mild (**B**) and severe (**C**) reduction in perfusion pressure, whereas small arteries tend to constrict. The gray box indicates vessel diameters in the Liao and Kuo dataset [5]. Literature data are presented as mean ± SD.

#### 3.3.3 Diameter responses

**Figure 10** shows the simulated subepi diameter changes during mild (*p*^*in*^ = 59 mmHg, panel B) and severe (*p*^*in*^ = 38 mmHg, panel C) reductions in coronary perfusion pressure using the low and high *ϕ*_*myo*_ profiles, compared with experimental data on epicardial canine vessels [49]. For a mild reduction in coronary perfusion pressure with the low *ϕ*_*myo*_ profile (**Figure 10B**), the subepi arterioles (*D*_*h*_ < 160 μm) dilate, while the subepi small arteries vasoconstrict. Under the severe perfusion pressure reduction (**Figure 10C**), subepi arterioles dilated similarly to the mild reduction, while small arteries exhibited even greater constriction (**Figure 10C**). The high *ϕ*_*myo*_ profile had minimal impact on arteriole response under both mild and severe pressure reductions, likely because these vessels were already nearly fully dilated (**Figure 9**). In contrast, small arteries showed markedly reduced vasoconstriction with the high *ϕ*_*myo*_ profile (**Figure 10B,C**). The general response of our coronary tree model agrees with the experimental trends presented in Kanatsuka et al. [49].

### 3.4 Assessing the relative contributions of the autoregulatory mechanisms

The partitioning of the autoregulatory stimuli in **Equation (6)** allows for the assessment of the relative contributions of each individual mechanism. In **Figure 11A**, we systematically remove autoregulatory stimuli while maintaining the basal homeostatic stimulus (*s*_*h*_) in all cases. The metabolic-only case (i.e. with *s*_*myo*_ = *s*_*τ*_ = 0) yields a greater autoregulatory response compared to the myogenic- and shear-dependent-only cases (**Figure 11A**). The myogenic-only case still maintains some autoregulatory capabilities, whereas the shear-dependent-only case results in a complete loss of coronary autoregulation. Furthermore, when only shear-dependent control is removed, the resulting changes in coronary autoregulation are minor, further emphasizes that shear-dependent control has minimal contributions to pressure-flow autoregulation. These results indicate that the relative strengths of the autoregulatory stimuli follow the order: metabolic > myogenic > shear-dependent. However, when replacing the low *ϕ*_*myo*_ with the high *ϕ*_*myo*_ distribution (**Figure 2B**), myogenic control substantially strengthens the pressure-flow autoregulatory response. Thus, these conclusions depend on our assumed normalized autoregulatory strength distributions.

**Figure 11:**
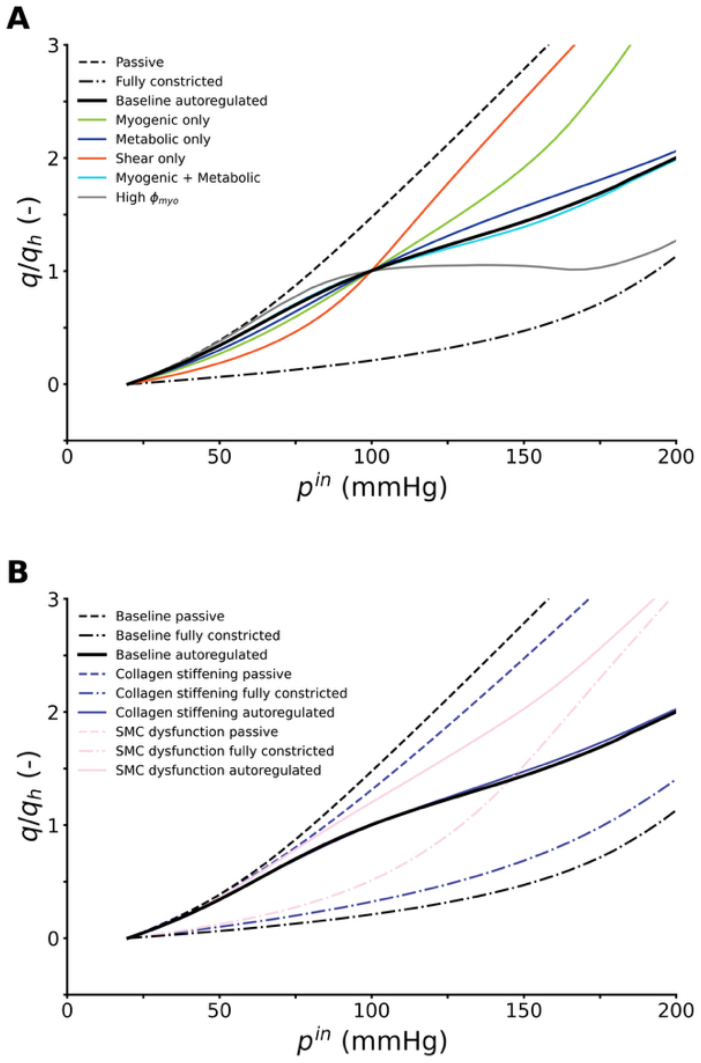
The impact of coronary pressure-flow autoregulation with (**A**) changes in autoregulatory stimuli and (**B**) change in vessel material properties.

### 3.5 Assessing the impact of vascular dysfunction on autoregulation

A key benefit of a microstructurally motivated model of coronary autoregulation is its integration of microstructural properties and cellular-level functions within the vessel wall model, enabling investigation into how structural and cellular changes affect coronary circulation function. We demonstrate the utility of our modeling framework by examining two scenarios: (1) stiffening of collagen fibers and (2) impaired SMC contractility. Specifically, collagen fibers are stiffened by doubling the passive material parameter *c*_2_, while SMC contractility is impaired by halving *S*_*max*_. The effects of these two scenarios on flow-pressure autoregulation are shown in **Figure 11B**.

Stiffening of the collagen fibers results in upward and downward shifts of the fully constricted and dilated curves, respectively (**Figure 11B**). However, coronary autoregulation remains relatively intact, suggesting that collagen stiffening constrains the dilatory capacity of the coronary tree without abolishing its autoregulatory function. In contrast, impaired SMC contractility produces a loss of coronary autoregulation accompanied by an upward shift in the fully constricted curve (**Figure 11B**). This indicates that reduced SMC contractile capacity severely compromises the ability of the coronary circulation to autoregulate.

## 4. DISCUSSION

The coronary circulation manifests a complex dynamic response to changes in perfusion pressure over multiple timescales. These timescales span from autoregulatory variations in active SMC tone, taking place within minutes, to slower changes in vessel microstructure [50], [51], [52] or vascular networks angiogenesis or rarefaction over the course weeks to months [53].

Recent computational modeling endeavors in coronary physiology focused on capturing key features of short-term coronary autoregulation using lumped-parameter models [11], [13], [14], [54]. However, because these approaches lack microstructurally-motivated models for vascular tissue, they are not equipped for predicting long-term pathophysiologic responses of the coronary circulation. To address this need, this work aimed to produce a unified computational approach that could integrate vascular responses at the short time scales (e.g., autoregulation) as well as long time scales (e.g. vascular growth and remodeling) while including morphometric, hemodynamics and structural data (**Figure 1A**). This design goal resulted in the adoption of a microstructurally-motivated modeling framework, widely used by the tissue growth and remodeling community, based on constrained mixture theory. The constrained mixture enables integration of microstructural properties and cellular level functions within the vessel wall in a nonlinear continuum mechanics framework. To our knowledge, this is the first implementation of a constrained mixture theory to model coronary autoregulation. The main findings of the model are discussed next.

### 4.1 Constrained mixture model parameter estimation

The vascular wall model was calibrated using passive and myogenic pressure-diameter data from *ex vivo* myography measurements in swine across four vessel sizes [5]. To apply this data to all myocardial layers, we assumed that the passive and myogenic response does vary transmurally across the myocardium for the transvascular pressures tested experimentally (*p*^*tv*^ ≤ 100 mmHg). To date, only a limited number of studies have investigated whether the myogenic response varies transmurally across the myocardium [42], [43], and it remains inconclusive whether meaningful differences exist [26].

Our calibrated model parameters include passive and active material properties, constituent pre-stretches, and mass fractions (**Table 2 and Table 3**). Our calibrated model effectively captures the passive and myogenic responses for all vessel sizes, at each myocardial depth (**Figure 5A-D**). For *p*^*tv*^ > 100 mmHg, we observe clear differences in the myogenic response between the three subtrees, with the subepi layer experiencing a greater myogenic response compared to the subendo layer. The myogenic response varies transmurally across the heart partly because blood vessels become thicker at greater depths in the myocardium (**Figure 6D**), which increases the total mass of load-bearing constituents. Future studies probing the myogenic response across a broader range of pressures would elucidate whether myogenic control varies throughout the myocardium.

Myocardial location is a determining factor of the microstructure of calibrated coronary vessels. SMC mass fraction is greatest in the subendo layer and lowest in the subepi layer for all vessel sizes (**Figure 5E**). The subendo layer likely has the greatest SMC mass fraction to compensate for its thinner vessels (**Figure 6D**), enabling it to generate sufficient active tension to capture the myogenic response. For each myocardial layer, our model estimates the same qualitative changes in mass fractions across the four vessel sizes (**Figure 5E**). Elastin mass fraction remained <10% for all vessel sizes considered, which aligns with our vessels being muscular in nature. To date, mass fractions of coronary artery constituents have only been quantified in large epicardial arteries [55], while the microcirculation remains understudied. Microscopic analysis of rabbit arterioles showed that SMC content gradually decreases from 100 *μ*m to 30 *μ*m vessels [56], consistent with our SMC mass fractions decreasing from intermediate to small arterioles. However, a quantitative assessment of the wall constituents was not performed. Future work quantifying the microstructural composition of the coronary microcirculation will enhance our understanding of coronary physiology and pathophysiology [57].

### 4.2 Morphometry, structure, and hemodynamics of homeostatic coronary trees

In stage two of model development, the homeostatic optimization method proposed by Gharahi et al. (2023) was used to generate three symmetrically bifurcating coronary trees in subepi, midwall, and subendo layers [24]. This approach estimates the homeostatic morphometric (e.g, diameters), structural (e.g, thicknesses), and hemodynamics (e.g., wall shear stress) characteristics of a vascular tree by minimizing the metabolic and viscous energy dissipation under the constraint of mechanical equilibrium. Furthermore, to incorporate measurements of the bifurcation diameter exponent (*ξ*)[46] into the homeostatic optimization, a penalty term was added into the cost function.

The estimated morphological and homeostatic characteristics are summarized in **Figure 6**. Swine measurements by Guo and Kassab (2004) [20] were used as targets for the subepi wall thickness, whereas the midwall and subendo thicknesses were calibrated to match the subepi hoop stress distribution. The decision to match the hoop stress distributions of the three subtrees was based on the experimental findings that vessel wall thickness varies transmurally across the myocardium [9], suggesting that mechanical stimuli play an important role in determining vessel thickness [9], [58].

In our framework, vessel thickness is calibrated by tuning SMC homeostatic activation (*A*_*h*_). The predicted homeostatic activation level decreases across generations 1-4 and 7-12, while generations 4-7 deviate from this trend (**Figure 6E**). The decreasing *A*_*h*_ across generations 1-4 and 7-12 is likely a consequence of assigning constant material properties to vessels outside the diameter range used for vessel wall model calibration (i.e., all small arteries and small arterioles have the same material parameters). The pressure-diameter data from Liao and Kuo (1997) encompasses vessel diameters of 35-180 μm [5], whereas our tree spans diameters 10-500 μm. Using variable material parameters might be more appropriate and aid in generating a vessel-size independent distribution of *A*_*h*_ [25], but we chose not to because our current model already contains many parameters and uses nonlinear constitutive models. Therefore, the distribution of *A*_*h*_ should be interpreted cautiously. Our model would benefit from further characterization of active and passive material properties beyond the experimental range utilized in this study.

Our pressure and flow distributions showed reasonable agreement with previous modeling efforts [45]. We must note that the inlet pressure for all subtrees were assumed to be 100 mmHg based on the study by Mittal et al. [59] that showed the pressure does not significantly drop in vessels of 100-1000 μm. Our wall shear stress tends to be greater than that measured by Stepp et al. (1999) in canines [32], but agrees better with previous modeling work by Cornelissen et al. (2002) [11]. Collectively, stage 2 calibration produced coronary trees at three myocardial depths which have morphological and hemodynamical characteristics which reasonably agree with those reported in literature (**Figure 6**).

### 4.3 Coronary autoregulation model

Our coronary autoregulation model is inspired by models previously proposed in the literature. The SMC activation model proposed by Carlson et al. (2008) [29] was used for autoregulatory control, which modulates SMC tone as a sigmoidal function. In addition, we formulated the autoregulatory stimuli in terms of their deviation from a homeostatic value, which was motivated by previous applications of constrained mixture models in the study of long-term active SMC-mediated adaptations in cerebral vasospasms [23].

Our model successfully captures pressure-flow coronary autoregulation over a pressure range of 80-140 mmHg (**Figure 7A**); however, it showed larger deviations from experimental data at lower coronary pressures (20-60 mmHg). Over this lower pressure range, our model closely follows the fully passive curve, with SMCs being nearly fully dilated (**Figure 9**). Since even the fully passive curve falls below the targeted flow levels (**Figure 7A**), our inability to capture the flow-pressure relationship between 20-60 mmHg appears related to the passive mechanics of our vessels. As previously mentioned, vessel passive and myogenic mechanics are determined by *ex vivo* myography data and therefore neglects the effect of myocardial tethering. Myocardial tether plays an important role constraining the vessels [13], [60], and therefore, will be important in capturing the *in vivo* pressure-diameter response. Future efforts should therefore attempt to include myocardial tethering effects in the vessel wall model.

We explored the response of our autoregulation model to changes in epicardial pressure. **Figure 10A** shows that reduced coronary perfusion pressure decreases the ENDO/EPI flow ratio, consistent with the known susceptibility of the subendo layer to ischemia [49], [61], [62]. Additionally, increases in perfusion pressure resulted in increased ENDO/EPI ratios, as reported by Boatwright et al. (1980) [10]. Thus, our model captures a transmural variability in autoregulatory capacity. Changes in ENDO/EPI are partially a consequence of subendo vessels becoming fully dilated or constricted within a narrower pressure range when compared to subepi vessels, illustrated by the steeper SMC activation curves in subendo vessels (**Figure 9G,N**). These results demonstrate by ability of our model to successfully integrate experimental morphometric and structural data to recapitulate key patterns of coronary autoregulation.

We also compared changes in vessel diameters in response to mild (*p*^*in*^ = 59 mmHg) and severe (*p*^*in*^ = 38 mmHg) reductions in coronary perfusion pressure with those reported by Kanatsuka et al. (1990) in canines [49]. We found that small arteries constricted while arterioles dilated, in agreement with experimental measurements (**Figure 10B,C**). However, our large vessels exhibited greater constriction compared to the experimental data. By increasing myogenic reactivity in small arteries, we were able to reduce the degree of constriction in these vessels (**Figure 10B**), leading to a better agreement with experiment. This suggests that vessels >200 μm may have higher myogenic strength than currently prescribed in our low *ϕ*_*myo*_ profile. Furthermore, arterioles dilated to a smaller degree than reported, which could not be remedied by increasing the myogenic strength due to vessels already being fully dilated (**Figure 9 and Figure 10B,C**).

The dilatory ability of the vessel is largely a consequence of the vessel wall calibration (stage 1). Through repeated testing, we found that we were unable to increase the dilatory ability of the vessels, while still matching both the passive and myogenic pressure-diameter data. For both the mild and severe pressure reduction, our model showed a maximum dilation occurring at a vessel diameter of 72 μm, and a continuously decreasing amount of dilation for smaller vessels. This behavior is also captured in the mathematical model developed by Cornelissen et al. (2002) in response to adenosine and L-NAME infusion in the coronary microcirculation [11], [63].

### 4.4 Model sensitivity

Through our sensitivity analysis, we identified small arteriole pre-stretches and *S*_*max*_ as the most sensitive parameters (**Table 2**: Constrained mixture model parameters that are constant across vessel sizes. **and Table 3**). This finding is physiologically reasonable, as small arteriole pre-stretches directly influence stiffness of vessels that provide the greatest resistance to coronary flow, while *S*_***max***_ plays a central role in determining the level of active tension. Note, our sensitivity analysis only considered the pressure-flow relationship (**Figure 7A**) and does not consider the passive and myogenic data (**Figure 5A-D**). Parameters may appear as not sensitive through this analysis in terms of impacting the pressure-flow relationship but could considerably impact the pressure-diameter relationship.

By knocking out each of the autoregulatory mechanisms, we were able to investigate their relative contributions (**Figure 11A**). We found that metabolic control to be the most dominant in pressure-flow autoregulation, followed by myogenic, and then shear-dependent (**Figure 11A**). Shear-dependent control was found to only provide a minor contribution, and removing it marginally impacted pressure-flow autoregulation, consistent with Namani et al. (2018) [13]. The qualitative contributions between the three autoregulatory mechanisms agree with Figure 2A of Carlson et al. (2008) [29] for arterioles supplying skeletal muscles. We found that increasing the normalized myogenic response in the small arterioles had a profound impact on the pressure-flow relationship, meaning the relative contribution of the myogenic response is strongly dependent on our normalized myogenic response distribution.

A key advantage of our coronary autoregulation model is its incorporation of microstructural properties and cellular-level function within individual vessels. We demonstrated the utility of this microstructurally-motivated framework by examining how collagen stiffening or reduced SMC contractility affects coronary pressure-flow autoregulation. We found that collagen stiffening had minimal impact on pressure-flow autoregulation but significantly reduced coronary flow reserve (**Figure 11B**). This suggests that while adequate coronary flow may be maintained under resting conditions, the stiffer vessels exhibit insufficient vasodilation during periods of increased metabolic demand, thereby compromising flow reserve capacity. Halving *S*_***max***_ resulted in complete loss of pressure-flow autoregulation, highlighting the critical role of SMC contractile function in maintaining coronary autoregulatory capacity.

### 4.5 Pulsatile coronary hemodynamics

Coronary flow simulations were performed by extending Womersley’s solution to account for an external time-varying intramyocardial pressure [38]. This approach allows our model to capture important coronary flow physics that are neglected in 0D lumped parameter models, including wave propagations and reflections that influence pressure and flow waveforms (**Figure 8**). Furthermore, each vessel has a longitudinal spatial variable, removing the spatial ambiguity inherent in lumped parameter models.

Flow is diastolically dominant for all vessels in the midwall and subendo layer, whereas flow is more evenly split between systole and diastole in the subepi layer (**Figure 8**). The subepi layer experiences the lowest *p*^*im*^, and thus, the lowest impedance due to vessel compression. Capturing pulsatile flow waveforms across the heart is important for understanding the spatial heterogeneity of myocardial perfusion and plays an important role in oxygen delivery [15]. Collectively, our results show that the magnitude and distribution of *p*^*im*^ are important influences on coronary flow waveforms across the myocardium.

### 6. Model limitations

The present study has several limitations. First, the experimental measurements delineating the pressure-diameter relationships of myocardial coronary vessels are scarce. We used four available sets of pressure-diameter data for a wide range of subepi, midwall, and subendo vessels from 500 μm to 10 μm. More pressure-diameter data in the coronary arterioles will enhance the accuracy and predictive capabilities of the model. In this work, coronary vascular networks were idealized to bifurcating trees with 12 generations, whereas a realistic reconstruction of the coronary network could impact the local hemodynamics and autoregulatory responses. Lastly, understanding the mechanisms involved in the coronary autoregulation, especially the role of metabolic control, has been a largely debated issue in coronary physiology [64]. In this study we simulated coronary autoregulation using the three main mechanisms and their range of influence as defined in previous modeling studies. Our model, however, could be further refined with improvements in our understanding of the interplay between the different mechanisms of coronary autoregulation.

## 5. CONCLUSION

The study of physiology and pathophysiology in coronary circulation benefits from accurate mathematical models that can account for both the microstructure and physiological function of arterial networks. In this study, we presented a microstructurally-motivated coronary autoregulation model that uses a nonlinear continuum mechanics approach to account for the morphometry and vessel wall composition in two idealized coronary trees. Literature data were used to calibrate and test our model. With some modifications, this model can be applied to morphometry-based coronary trees instead of idealized symmetric trees. Finally, since our model is based on constrained mixture theory, it could be expanded to also study long-term growth and remodeling in the coronary circulation in response to hypertension, atherosclerosis, etc.

## Nomenclature

a: Autoregulatory scaling coefficient
A: SMC activation
c: Pulse wave velocity
c_1_: Elastin material parameter
c_2_: Collagen material parameter
c_3_: Collagen dimensionless material parameter
c_4_: SMCs material parameter
c_5_: SMCs dimensionless material parameter
CEP: Cavity-induced pressure
D: Internal vessel diameter
G: Tissue constituent pre-stretch
H: Wall thickness
i: 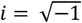
J: Bessel functions
L: Vessel length
L_0_: Metabolic characteristic length
L_p_: Path length
LV: Left ventricle
M: Mass per unit reference area of a tissue constituent
MVO_2_: Myocardial oxygen consumption
NO: Nitric oxide
p: Lumen pressure
p^im^: Intramyocardial pressure
p^LV^: Left ventricle pressure
p^tv^: Transvascular pressure
q: Flow rate
s: Autoregulatory stimulus
S_max_: Maximum SMC activation stress
SIP: Shortening-induced pressure
SMC: Smooth muscle cells
T_θθ_: Circumferential wall tension
w: Strain energy density
Y: Vessel admittance
z: Longitudinal spatial coordinate
α_Wom_: Womersley number
α_myo_: Myogenic scaling coefficient
α_meta_: Metabolic scaling coefficient
α_τ_: Shear-dependent scaling coefficient
β: Normalized myocardial depth
θ: Circumferential spatial coordinate
λ_0_: Zero stretch
λ_M_: Maximum active tension stretches
λ_z_: Axial stretch
λ_θ_: Circumferential stretch
μ: Blood viscosity
v: Tissue constituent mass fraction
ξ: Bifurcation diameter exponent
ρ_blood_: Blood density
ρ_wall_: Density of the vascular wall
σ: Hoop stress
τ: Wall shear stress
ϕ: Normalized autoregulatory response
ω: Angular frequency

## ACKNOWLEDGEMENTS

This study was supported by the NIH R01-HL158723, and U01-HL135842, R01-HL139813.

## Appendix A Mechanics of a single artery

A single vessel of the arterial tree is considered as a thin-walled cylindrical tube composed of three main load-bearing constituents: elastin (*e*), collagen (*c*), and smooth muscle cells (SMCs). First, we only consider the passive response of constituents. Each constituent is assumed to separately contribute to the strain energy density:

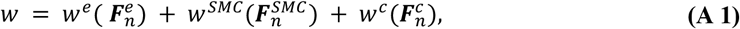

where 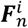 is the deformation gradient of each constituent *i* (*i* ∈ {*e, SMC, c*}) corresponding to its map from a stress-free configuration to the overall homeostatic configuration. We define this deformation gradient as 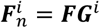, where ***F*** represents the deformation gradient of the mixture mapping from its reference configuration to the homeostatic configuration, and ***G***^*i*^ is a pre-stretch for each constituent mapping each constituent from its distinct stress-free configuration to the reference configuration [65]. In particular, the pre-stretch mapping for elastin can be expressed as

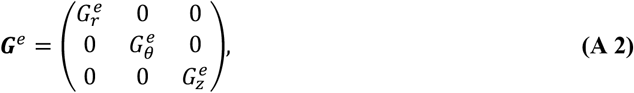

where 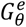 and 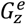 are pre-stretches associated with circumferential and axial directions, and 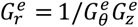.

Similarly, for collagen fibers and SMCs, ***M***^***i***^, *i* ∈ {*k, SMC*} is defined as the unit vector in the direction of the collagen fiber (*k*) or SMCs. The pre-stretch mappings for collagen and smooth muscle cells are given as

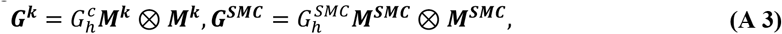

where the pre-stretches 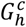 and 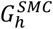 are also called homeostatic stretches, the stretches of the constituents when they are produced. We should note that in the previous applications of growth and remodeling, the pre-stretches were assumed to be constant for a single vessel. In our generalization of the framework to a vascular tree, we account for the variation of pre-stretches across the generations of vessels. Nevertheless, the pre-stretch implies that the homeostatic state in an individual vessel is associated with a constant homeostatic stress for the constituents of the vessel wall.

The orientation of collagen fibers and smooth muscles with respect to the axial direction in their reference configuration, defined by angle *γ*^*k*^, can be written as

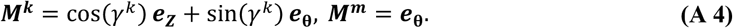

For modeling the extension and inflation of a thin wall model, ***F*** is considered as ***F*** = *diag*[*λ*_*r*_, *λ*_*θ*_, *λ*_*z*_]. The stretches of constituent *i, λ*^*i*^, *i* ∈ {*e, k, SMC*}, are expressed in terms of the pre-stretches using ***F***^*i*^ = ***FG***^***i***^

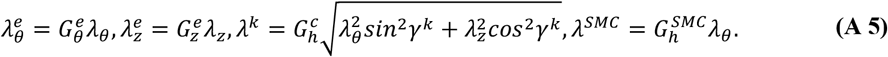

The incompressibility of the wall material is imposed by assuming an isochoric motion (i.e., det(***F***) = 1), and thus *λ*_*r*_ = 1/*λ*_*θ*_*λ*_*z*_. Using the membrane theory [28], the passive membrane Cauchy stress (force per deformed length) can be written as

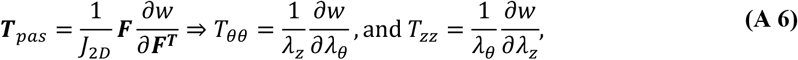

where *J*_2*D*_ = *λ*_*θ*_*λ*_*z*_. The total strain energy per unit area can be written as

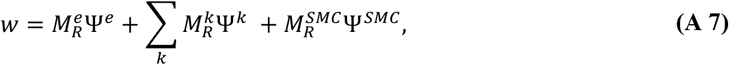

where 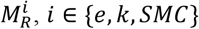 is the mass of each constituent per unit reference area. Alternatively, the total strain energy per unit area can be written as

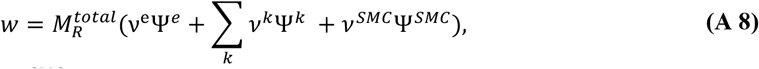

where *ν*^e^, *v*^*k*^, and *v*^*SMC*^ are mass fractions of elastin, collagen fiber families, and SMCs, respectively. In this work, four families of collagen fibers in circumferential, axial, and two diagonal directions were considered with mass fractions *v*^*k*^ =(0.1, 0.1, 0.4, 0.4)*v*^*c*^ where *v*^*c*^ is the total collagen mass fraction [66]. The total mass per unit area 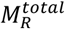 is the mass of load bearing constituents and can be computed via

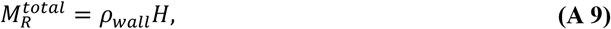

where *ρ*_*wall*_ is the density of the vascular wall and *H* is the thickness under homeostatic conditions. A neo-Hookean model is employed for the passive elastin response and a Holzapfel exponential model is used for collagen fiber families and passive behavior of circumferentially oriented SMCs

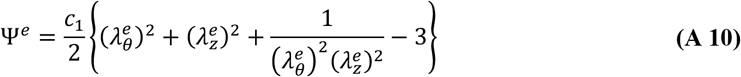

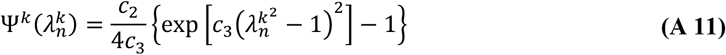

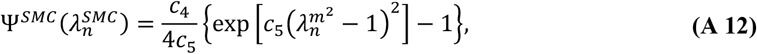

where *c*_1_ is the elastin material parameter, *c*_2_ and *c*_3_ are collagen material parameters, and *c*_4_ and *c*_5_ are passive SMC material parameters. To include the active tension of vascular SMCs, we use a potential function as given by [23]:

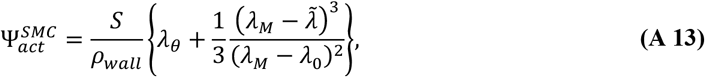

where *λ*_*M*_ and *λ*_0_ are stretches at which the active force generation is maximum and zero, respectively, and *ρ*_*wall*_ is the vascular wall density. In addition, 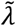 is an active stretch of the SMCs in the circumferential direction, which can evolve by SMC remodeling over slow timescales (hours to days). In the current study, we focus on the short timescale adaptations (minutes), and thus, 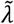 is assumed to be the total circumferential stretch in the vessel (*λ*_*θ*_). Active SMC stress *S* is in general a function of the myogenic, shear-dependent, and metabolic controls in coronary vessels. Again, using membrane theory, 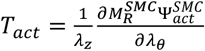 the total tension in the artery can be written as

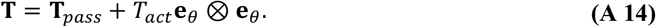

Finally, for a thin-walled cylinder with transmural pressure *p*^*tv*^, the force equilibrium in the circumferential direction gives

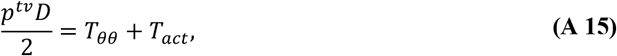

where *D* is the vessel diameter and *p*^*tv*^ is the transvascular pressure of the artery.

We assume the reference configuration of the blood vessel in our continuum mechanics formulation is its homeostatic configuration. Therefore, by setting ***F*** = ***I*** (i.e., *λ*_*θ*_ = *λ*_*z*_ = 1 ) in equations above, we can write

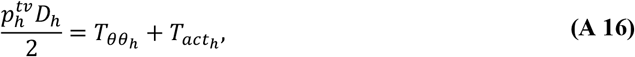

where 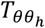 and 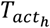 are homeostatic passive and active circumferential tensions, respectively, 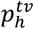 is the homeostatic transvascular pressure and *D*_*h*_ is the homeostatic diameter of the vessel. The homeostatic passive tension 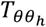 is thus determined by the material properties (*c*_1_-*c*_5_), constituent prestretches (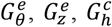, and 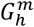) and their mass fractions *v*^*i*^. In addition, the homeostatic active tension 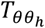 is determined by active SMC parameters, *λ*_*M*_, *λ*_0_ and *S*_*h*_. Accordingly, we can also compute the homeostatic stress in each constituent ***σ***^***i***^ as a function of its material parameters, its homeostatic stretch *G*^*i*^, and active SMC stress in case of SMCs.

## Appendix B Womersley solution admittance matrix

At a given non-zero frequency *ω*, the Womersley’s solution for transvascular pressure and flow are calculated as [37], [38]:

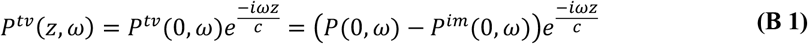

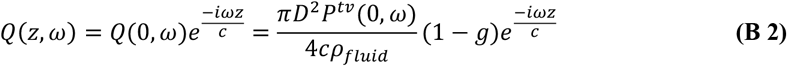

where *P*(0, *ω*) is the inlet lumen pressure, *P*^*im*^(0, *ω*) is the inlet intramyocardial pressure, *P*^*tv*^(0, *ω*) is the inlet transvascular pressure, and *Q*(0, *ω*) is the inlet flow, all in the frequency domain. The variable *z* is the longitudinal spatial coordinate, *c* is the pulse wave velocity, *ρ*_*blood*_ is the blood density, 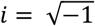, and *g* = 2*J*_1_(∧)/∧ *J*_0_(∧). *J*_0_ and *J*_1_ are Bessel functions of the first kind, and 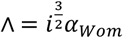 with *α*_*Wom*_ being the Womersley number defined as 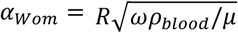 and *μ* being the blood viscosity. Utilizing Euler’s formula, **Equation B 1** and **B 2** can be rewritten as:

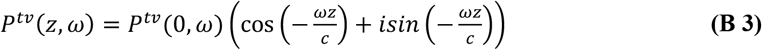

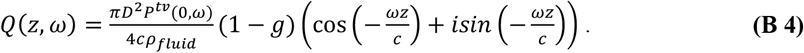

For a vessel of length *L*, we wish to solve for the flow, provided pulsatile pressure boundary conditions. For notational simplicity, let 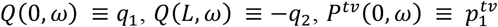, and 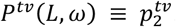. We seek an expression of the form: ***q*** = ***Yp***^***tv***^, where ***Y*** is an admittance matrix, ***q*** = [*q*_1_ *q*_2_]^*T*^ and 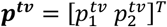. To solve for ***p***^***tv***^ and ***q***, we plug *z* = 0 and *z* = *L* into **Equation B 3** and **B 4**. This results in:

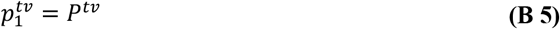

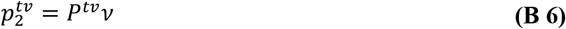

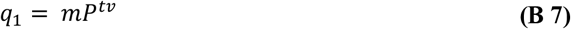

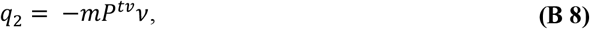

With 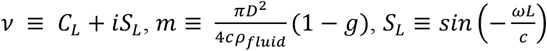, and 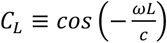. **Equations B 5-8** are written as a system of equations:

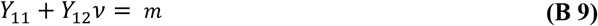

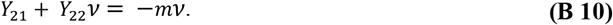

Furthermore, we have *Y*_11_ = *Y*_22_ and *Y*_12_ = *Y*_21_ due to the symmetric nature of the system. This results in the admittance matrix:

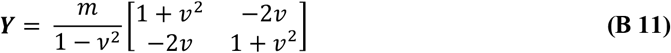

## Appendix C Viscosity

Blood viscosity in the systemic vasculature depends on the size of the vessel and the hematocrit level (*H*_*D*_). Particularly, the variation of viscosity is more pronounced as the arteries and arterioles become smaller. In our study, we prescribe the viscosity using the following *in-vivo* viscosity law given in [39]:

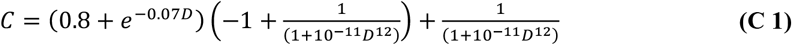

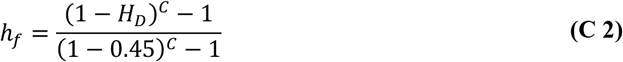

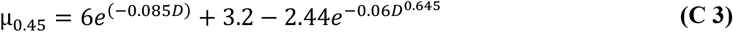

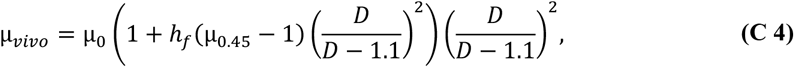

where *D* is the diameter in microns, and μ_0_ is the viscosity of the blood plasma taken to be 0.001 Pa.s.

## Appendix D CMM parameter estimation procedure

Due to the large number of CMM parameters, Stage 1 calibration is performed iteratively. Parameters are classified as either global or local. Global parameters do not dependent on vessel size or myocardial depth and include the material constants (*c*_1_ − *c*_5_) and SMC active parameters *S*_*max*_, *λ*_0_, and *λ*_*M*_. Note, SMC active material parameters are determined through literature values **Table 2**. Local parameters are those which vary between the vessel sizes and include all pre-stretches (*G*), mass fractions (*υ*^*e*^ and *υ*^*c*^), and myogenic strengths (*a*_*myo*_ and *ϕ*_*myo*_). Note, SMC mass fraction is determined as: *υ*^*m*^ = 1 − *υ*^*e*^ − *υ*^*c*^.

The calibration procedure is performed by iteratively optimizing global and local parameters. For optimizing global parameters, local parameters are fixed, and global parameters are calibrated using Nelder-Mead optimization [67]. To calculate the global error, the local error of each individual vessel, as defined in **Equation (20)**, is summed. For optimizing local parameters, global parameters are fixed, and local parameters are calibrated over each individual vessel. We iterate between global and local parameter optimization until our parameters converge.

## Appendix E Homeostatic optimization

Originally introduced by Murray [68], and later extended by Taber [69] and Lindström et al. [70], the extended Murray’s law states that the vessel wall composition and geometry strive to minimize the energy consumption. The homeostatic optimization framework proposed in [24] leverages this idea to identify the homeostatic characteristics of an arterial tree. Briefly, the total energy cost per unit length *C* for an individual blood vessel can be written as:

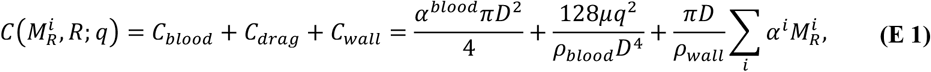

where *C*_*blood*_ is the metabolic cost of sustaining blood in the vessels, *C*_*drag*_ is the power needed to overcome viscous drag, and *C*_*wall*_ is the metabolic cost of maintaining the constituents of the wall. In addition, *α*^*blood*^ and *α*^*i*^ are the metabolic energy costs of blood and wall constituents (*i* ∈ {*e, k, SMC*}, see **Appendix A**) per unit volume, *q* is the blood flow in the vessel, and *μ, ρ*_*blood*_, and *ρ*_*wall*_ are the blood viscosity, blood density, vessel wall density, respectively. We must note that the metabolic cost of SMCs can be written as the summation of the metabolic cost of their maintenance 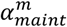 and the metabolic cost of the active SMC stress 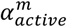.

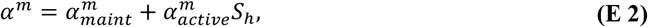

where *S*_*h*_ is the homeostatic active SMC stress. This metabolic cost function must be minimized with Eq. A.14 as the constraint in each segment. With regards to Appendix A, we can rewrite the mass of constituents in terms of their mass fractions as 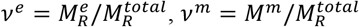, and 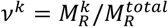 with *v*^*c*^ = ∑*v*^*k*^. Using the mass fractions and considering that the arteries are in homeostatic state, we can write the mechanical equilibrium in each vessel as

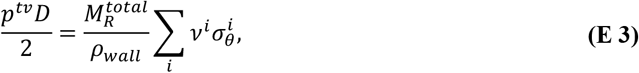

where 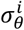 is the circumferentially acting part of the homeostatic stress tensor ***σ***^***i***^. Therefore, the energy cost of one vessel can be written in terms of diameter, with pressure and flow rate given from the hemodynamics as

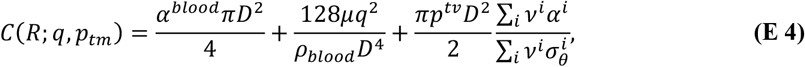

The total energy cost of all arteries can be added to compute the total cost for maintenance of an arterial tree

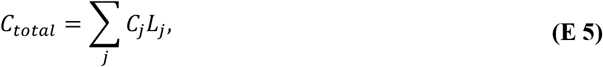

where *C*_*j*_ is the energy cost (Eq. B.3) and *L*_*j*_ is the length of segment *j*. In this study, we add the bifurcation diameter exponent (*ξ*) measurement from Kaimovitz et al. [46] the pressures at the inlet and outlets and incorporate the flow rate as a penalty term to the cost function.

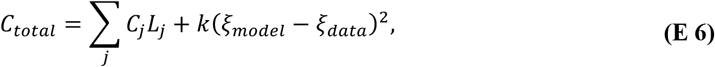

where *ξ*_*model*_ is the bifurcation diameter exponent in the tree, *ξ*_*data*_ is the measured bifurcation diameter exponent, and *k* is a penalty coefficient. Once the radius of each vessel is determined by minimization of *C*_*total*_, the thickness of each vessel can be computed using equations B.2 and A.10. As explained in **Appendix A**, the homeostatic stress tensor ***σ***^***i***^ for each constituent is a function of its material parameters, its homeostatic pre-stretches, and the active stress *S* for SMCs. In this work, to find the homeostatic characteristics of the vessel wall in the tree, material parameters and pre-stretches are determined according to their diameter using the 4 types of vessels; small arteries, large, intermediate, and small arterioles. However, the SMC active stress *S* is tuned for each individual vessel in the tree by changing SMC activation *A* to fit the thickness to diameter ratio reported in Guo and Kassab [20]. The parameters used for the homeostatic optimization are summarized in Table B1. thickness-to-diameter ratio with data in [20], as demonstrated in the results section.

**Table B1:** Parameters of the homeostatic optimization.

| Parameter | Description | Value | Unit | Reference |
| --- | --- | --- | --- | --- |
| $\alpha^c$ and $\alpha_{maint}^m$ | Metabolic cost of collagen and SMCs per unit volume | 1070 | W/m<sup>3</sup> | [71] |
| $\alpha^e$ | Metabolic cost of elastin per unit volume | 0 (Only produced at early ages.) | W/m<sup>3</sup> | [70] |
| $\alpha^{blood}$ | Metabolic cost of blood per unit volume | 51.7 | W/m<sup>3</sup> | [72] |
| $\alpha_{active}^m$ | Metabolic cost of active SMC stress | 0.00872 | 1/s | [71] |
| $\rho_{wall}, \rho_{blood}$ | Vessel and blood density | 1060 | kg/m<sup>3</sup> | - |

